# Rep virulence genes have been recruited convergently by nudivirus-derived Virus-Like Particles and ichnoviruses of Campoplegine wasps

**DOI:** 10.1101/2023.12.12.569846

**Authors:** Alexandra Cerqueira de Araujo, Matthieu Leobold, Marie Cariou, Bernardo Ferreira dos Santos, Rustem Uzbekov, Renato Ricciardi, Pierluigi Scaramozzino, Andrea Lucchi, Karine Musset, Jean-Michel Drezen, Thibaut Josse, Elisabeth Huguet

**Affiliations:** CBER, Department of Genetics, Evolution and Environment, University College London, London, UK; Institut de Recherche sur la Biologie de l’Insecte (IRBI), UMR 7261, CNRS - Université de Tours, Tours, France; Service Analyse de Données, Muséum National d’Histoire Naturelle, Centre National de la Recherche Scientifique, Paris, France; Museum für Naturkunde, Leibniz Institute for Evolution and Biodiversity Science, Center for Integrative Biodiversity Discovery, Invalidenstraße 43, Berlin, 10115, Germany; Université de Tours, Département des Microscopies, Tours, France; Faculty of Bioengineering and Bioinformatics, Moscow State University, Moscow, Russia; Department of Agriculture, Food and Environment (DAFE), University of Pisa, Italy

**Keywords:** Nudivirus, Viral domestication, Campopleginae, Virus-Like-Particle, Parasitoid wasp, Ichnovirus

## Abstract

Viral endogenization events in the genomes of parasitoid wasps have led to the production of immunosuppressive particles that are injected into the host and that either contain DNA encoding virulence effectors or virulence proteins, essential for parasitism success. In some Campopleginae (Ichneumonidae) wasps, viral endogenisation involves the production of large dsDNA ichnoviruses encoding several dozens of rep virulence genes, to the exception of *Venturia canescens*. Instead, *V. canescens* harbours an endogenous nudivirus producing Virus-Like-Particles (VLPs) in which virulence proteins are wrapped into viral envelopes. Here, we described that this nudiviral domestication event is more ancient and widespread than previously thought as we identified homologous alphanudivirus sequences in the genomes of three Campopleginae species belonging to the genera *Campoplex* and *Rimphoctona*. We showed that *Campoplex capitator* produces morphologically similar VLPs to those of *V. canescens* with almost the same viral protein content. Strikingly, the virulence proteins are however different, suggesting that VLPs allow a high versatility in the molecular strategies used by the wasps in response to host selection pressures. Furthermore, a marked expansion of serpin and *rep* genes is present in *Campoplex capitator*, with one serpin and several REP proteins identified as major VLP components using proteomics. REP proteins are also well-described virulence factors initially shown to be delivered into hosts via the expression of *rep* genes coded by ichnovirus DNA. In *Campoplex capitator*, these proteins are packaged into nudivirus-derived VLPs indicating that convergent use of virulence factors can imply endogenous viruses of different origin and with different delivery mechanisms.

## Introduction

The endogenization of viral sequences within eukaryotic genomes is a common phenomenon in the evolutionary history of eukaryotic organisms (Holmes 2011; Feschotte and Gilbert 2012). These viral sequences, known as endogenous viral elements (EVEs), involve all virus types likely to integrate into any eukaryotic genome (Bejarano et al. 1996; Delaroque et al. 1999; Crochu et al. 2004; Maori et al. 2007; Belyi et al. 2010; Katzourakis and Gifford 2010; Liu et al. 2010; Feschotte and Gilbert 2012; Ballinger et al. 2014; Irwin et al. 2021). While a large proportion of EVEs are counter-selected and no longer functional (Tarlinton et al. 2006; Arbuckle et al. 2010; Katzourakis and Gifford 2010), there are, however, examples of viral domestications providing novel adaptive functions and a selective advantage to the eukaryotic host, such as EVEs involved in placentation in mammals (Mangeney et al. 2007; Lavialle et al. 2013).

In general, this type of domestication involves individual viral genes rather than entire viral genomes. However, in some parasitoid wasp species, entire portions of large double-stranded DNA (dsDNA) viruses have been integrated and perform crucial functions in the wasp’s life cycle. Indeed, the wasps use the virus-derived particles produced in their ovaries and injected at the same time as wasp eggs in the insect host, to deliver virulence genes or virulence proteins that suppress the host immune response and spark other physiological changes to ensure parasitism success (Bézier, Herbinière, et al. 2009; Volkoff et al. 2010; Strand and Burke 2012; Burke et al. 2013; Strand and Burke 2014; Drezen et al. 2017).

To date, several viral domestication events have been identified in independent wasp lineages, each involving large dsDNA viruses (Burke 2019; Di Giovanni et al. 2020; Burke et al. 2021; Drezen et al. 2022; Guinet et al. 2023). These endogenization events can be classified into different categories based on the nature of the virus that has been endogenized, the wasp families involved, and the type of particles produced and their content (dsDNA circles containing virulence genes or virulence proteins).

Approximately 100 million years ago, the integration of sequences from a deltanudivirus (Petersen et al. 2022) in the ancestor of Braconidae Microgastrinae wasps – comprising at least 46000 species – led this lineage to produce Bracovirus (BV) particles, which are composed of a viral envelope enclosing capsids containing dsDNA circular molecules harbouring virulence genes (Murphy et al. 2008; Bézier, Annaheim, et al. 2009; Thézé et al. 2011; Herniou et al. 2013).

Two independent endogenization events leading to the production of Ichnovirus (IV) particles also containing circular dsDNA have involved a viral ancestor from a still uncharacterized virus family in two distinct Ichneumonidae lineages (Banchinae and Campopleginae) (Volkoff et al. 2010; Gundersen-Rindal et al. 2013; Strand and Burke 2014; Béliveau et al. 2015; Volkoff and Cusson 2020). The production of these particles involves the expression of several intronless genes tightly clustered into wasp genomic regions called IVSPERs (Ichnovirus Structural Protein Encoding Regions) (Volkoff et al. 2010; Volkoff et al. 2012; Lorenzi et al. 2019). Although Bracoviruses and Ichnoviruses do not have the same viral origin they share organisational and functional features, showcasing a remarkable example of convergent evolution. Indeed, in both cases virus particles are produced by a conserved set of endogenized viral genes of known (BV) or unknown (IV) origin. The particles deliver virulence genes in the parasitized host that are harboured by dsDNA circles, originating from linear segments in the wasp genome, that are circularized during BV or IV production thanks to conserved direct repeat junctions (Pasquier-Barre et al. 2002; Annaheim and Lanzrein 2007; Desjardins et al. 2008; Beck et al. 2011; Bézier et al. 2013). These BV and IV circles also have the capacity to integrate into the DNA of the parasitized insect host genomes, thanks to Host Integration Motifs present in some circles (Beck et al. 2011; Chevignon et al. 2018; Muller et al. 2021; Heisserer et al. 2023; Kim et al. 2023). The integrative property of these ds DNA BV and IV circles is most probably in large part at the origin of HGT events observed in the genomes of Lepidopteran species that are the main order targeted by these endoparasitoid wasps (Gasmi et al. 2015; Heisserer et al. 2023). The only similar virulence genes between BV and IV, possibly of wasp origin and likely resulting from horizontal transfer between these viruses, correspond to genes encoding ANK proteins involved in host immune suppression (Cerqueira de Araujo, Josse, et al. 2022). The other virulence genes, probably also of eucaryote origin, are not shared between BV and IV. Campopleginae-associated IV all contain a multigene virulence family having undergone particularly high expansion corresponding to repeat element genes (rep) (Volkoff et al. 2002; Tanaka et al. 2007; Legeai et al. 2020; Kim et al. 2023). Although the function of rep genes has not yet been elucidated, their conservation and high expansion in these IV genomes together with their reported expression in caterpillar hosts suggest that they play an important role in the parasitoid-host interaction (Galibert et al. 2006; Rasoolizadeh et al. 2009; Etebari et al. 2011; Kim et al. 2023).

Remarkably, nudiviruses have been involved in other independent domestications, with the endogenization of alphanudiviruses in the braconid *Fopius arisanus* (Burke et al. 2018) and more surprisingly in the Campopleginae *Venturia canescens* (Pichon et al. 2015). In these parasitoid wasps, the nudivirus endogenization event took a different evolutionary trajectory and has led to the production of Virus-Like-Particles (VLPs) that do not contain DNA but virulence proteins instead (Pichon et al. 2015; Burke et al. 2018; Burke 2019). Because *V. canescens* belongs to the subfamily Campopleginae, known to harbour ichnoviruses, the VLPs were initially thought to derive from IVSPER expression (Reineke et al. 2006). However, sequencing of the wasp’s genome identified only a few pseudogenised IVSPER genes, and it was formally demonstrated that the VLPs are in fact of nudiviral origin (Pichon et al. 2015). Venturia canescens Endogenous Nudivirus (VcENV), allows the production VLPs, which consist of a viral envelope containing virulence proteins that protect the wasp eggs from the lepidopteran host immune system after oviposition (Salt 1965; Rotheram 1967; Rotheram 1973; Bedwin 1979; Feddersen et al. 1986; Reineke et al. 2006; Pichon et al. 2015). Three main virulence proteins have been described in VcVLPs: *vlp1*, *vlp2*, and *vlp3* which encode respectively a glutathione peroxidase (PHGPx), a Rho GTPase-activating protein (RhoGAP), and a metalloprotease (neprilysin) (Theopold et al. 1994; Hellers et al. 1996; Asgari et al. 2002; Reineke et al. 2002; Pichon et al. 2015).

Several lines of evidence had suggested that the nudivirus integration event in *V. canescens* is more recent than the one described in the ancient braconid wasp 100 million years ago (Murphy et al. 2008; Thézé et al. 2011; Pichon et al. 2015). Indeed, to date, an alphanudivirus endogenization had only been described in a single campoplegine wasp species. Also, the nudivirus genes are less dispersed in the *V. canescens* genome than in braconid wasps (Bézier, Annaheim, et al. 2009; Burke et al. 2014; Pichon et al. 2015; Mao et al. 2023), suggesting fewer genomic rearrangements had time to occur. It has also been shown that in VcENV, losses of certain core nudivirus functions (*i.e*.: proteins involved in capsid formation) are due to pseudogenization involving accumulation of mutations rather than complete loss of genes or genomic regions as described for obligate symbionts of bacterial origin (Leobold et al. 2018). The fact that pseudogenised nudiviral sequences are not completely eroded could also be an indication that the nudivirus integration is more recent in this species (Leobold et al. 2018).

Here, we describe that this nudiviral domestication event is in fact more ancient than previously thought as we identified homologous alphanudivirus genomes in 2 species of the sister genus *Campoplex* by genome sequencing, and in *Rimphoctona* by data mining. We showed that *Campoplex capitator*, the main parasitoid of the vineyard pest *Lobesia botrana*, produces morphologically similar VLPs to those of *V. canescens* with almost the same viral protein content. Strikingly, the virulence proteins are completely different, suggesting that VLPs allow a high versatility in the molecular strategies used by the wasps in response to host selection pressures. Furthermore, a marked expansion of serpin and rep genes is present in *Campoplex capitator* genome, with one serpin and several rep proteins identified as major VLP components in proteomic analyses. Rep proteins are important virulence factors in wasps carrying ichnoviruses. They are produced from *rep* genes located on ichnovirus DNA circles once these circles have been delivered to the host. In *Campoplex capitator*, these virulence proteins are packaged into nudivirus-derived VLPs indicating that convergent use of virulence factors can imply endogenous viruses of different origin and with different delivery mechanisms.

## Results

### Viral integration Campopleginae wasps

In Campopleginae wasps, most species (of genera such as *Hyposoter*, *Campoletis*, *Diadegma*) have endogenized an ichnovirus and produce ichnovirus particles, while *Venturia canescens* was, so far, the only campoplegine wasp harboring an endogenized nudivirus and producing VLPs lacking capsids and DNA. To investigate whether closely related species possess endogenized viruses and determine their viral origin, we sequenced the genomes of two *Campoplex* species, belonging to a sister group of *V. canescens*: *Campoplex capitator* and *Campoplex nolae* (see Figure 1). Genome assemblies were obtained using hybrid sequencing approaches, yielding a 261 Mb assembly for *C. capitator* (630 contigs, N50 = 7.8 Mb, 96% BUSCO completeness, generated by PacBio sequencing) and a 218 Mb assembly for *C. nolae* (4,593 contigs, N50 = 1.8 Mb, 94% BUSCO completeness, generated by Illumina and Nanopore sequencing), with 11,288 and 13,929 predicted gene models, respectively (see table S1 for details). We have also extended our analyses to *Rhimphoctona megacephalus*, the most divergent species within the “Campoplex-Venturia-Dusona” clade after *Dusona sp.*, a species that does not harbour any confirmed endogenized virus (Burke et al. 2021) nor functional *rep* genes (see Figure 1).

**Figure 1:**
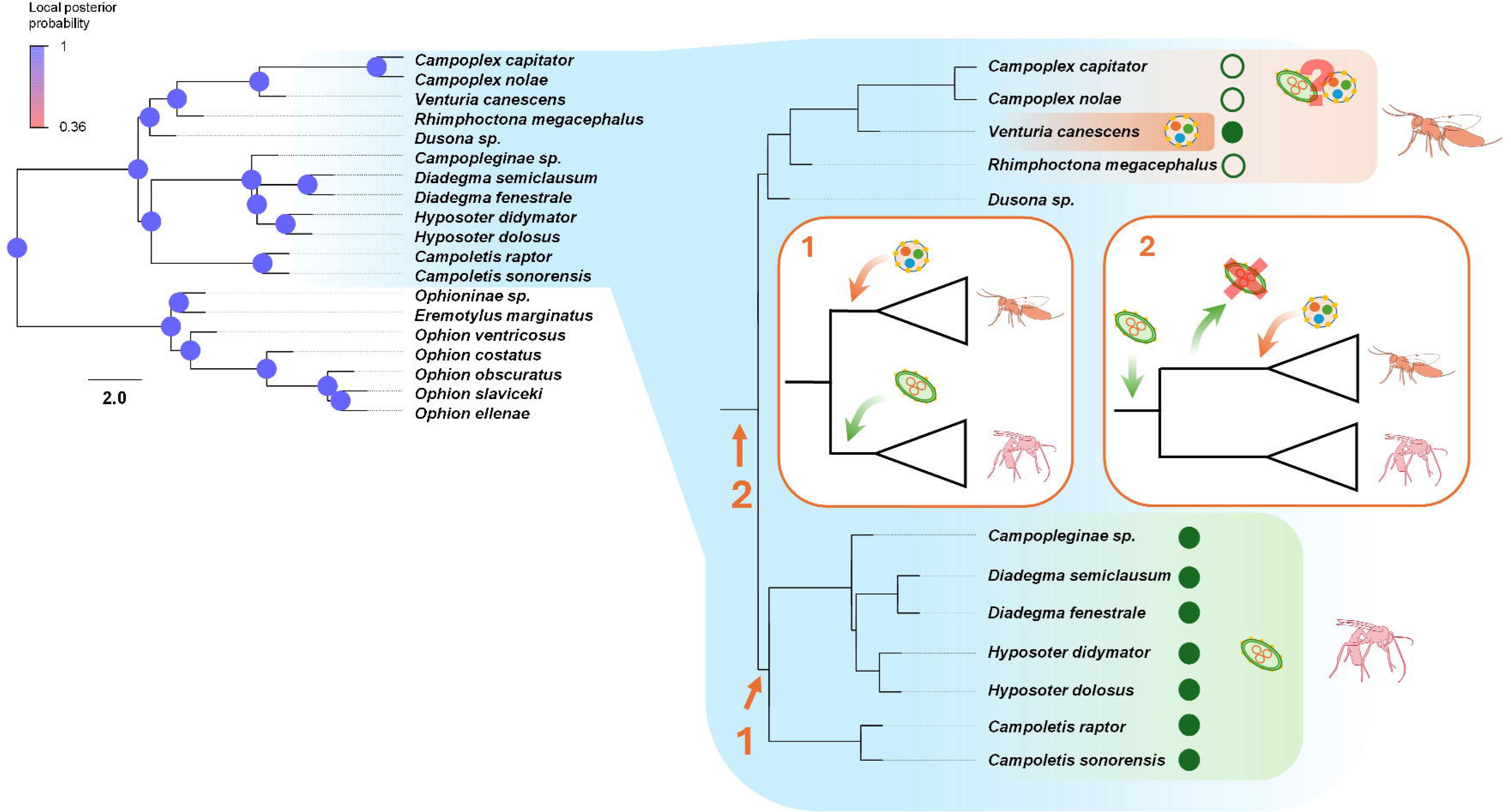
History of virus integration in Campopleginae. The left tree represents the Campopleginae and Ophioninae phylogenetic tree, built using hymenopteran BUSCOs extracted from NCBI genome assemblies. Branch topology has been inferred by coalescent modelling (ASTRAL) while branch lengths were inferred using a maximum likelihood approach. Branch support is given by local posterior probabilities, i.e. the probability that a branch is the true branch given the set of trees built using BUSCOs (computed based on the quartet score (Sayyari and Mirarab 2016)). A more complete tree of ichneumonid wasps is available in Figure S1. The right tree represents a portion of the Ichneumonidae tree corresponding to Campopleginae wasps. Solid green dots indicate species for which *rep* genes were found, while empty green circles indicate that *rep* gene presence needs to be investigated. We can distinguish two clades: one with an integrated ichnovirus (bottom part, in green), and one with *Venturia canescens*, which bears an integrated nudivirus (top part, in darker orange). The ichnovirus endogenized in *Campopleginae sp.* genome can be found in Guinet et al. 2023. Species closely related to *V. canescens* for which we do not know whether they bear a virus in their genome are mentioned in orange. So far, no viral integration has been observed in *Dusona spp.*. Two scenarios explaining viral integrations in Campopleginae can be envisaged: 1- The ichnoviruses and nudiviruses were acquired in two different branches of the Campopleginae tree. In this scenario, the nudiviral and ichnoviral sequences should not be found together and the two viruses did not interact with each other after integration. 2- An initial integration of ichnovirus took place in the common ancestor of all these species, followed by the integration of a nudivirus and certain branches of the tree. In this scenario, the ichnovirus could have been replaced by the nudivirus as a result of selection, and it would be possible to find both ichnoviral and nudiviral sequences, provided that the non-conserved sequences can still be characterised.

### Identification of endogenized nudivirus genomes within other Campopleginae wasp species : in *Campoplex spp.* and *Rimphoctona megacephalus* genomes

We identified 47 genes of viral origin in both *Campoplex* genomes and 62 viral genes in *R. megacephalus* genome (Table1 and Table S2). In *C. capitator*, these genes are almost exclusively expressed in adult wasp ovaries (Figure 2B and Table S3). Out of the 32 nudivirus core genes, 22 have been found in the *Campoplex* genomes and 21 in *R. megacephalus* (Table 1). Among these genes, some encode key viral functions in baculoviruses such as viral transcription (*p47* and *lef* genes), DNA amplification (*helicase*), envelope composition/infectiosity (*Ac81, p33* and *pif* genes) and virion morphogenesis (*vlf-1*). These genes are grouped into 4 clusters in *Campoplex spp.* and *R. megacephalus* genomes (Figure 2A for *C. capitator* and Figure S3A for *C. nolae*), but genes are more physically distant from each other in the *R. megacephalus* genome (see Figure S3B). In *V. canescens*, remnants of some nudivirus genes, notably those involved in DNA amplification and capsid formation, have been found as pseudogenes (Leobold et al. 2018). Therefore, for each missing nudivirus gene, we searched for remnant sequences in *Campoplex* and *Rhimphoctona* genomes. Several pseudogenes have been identified, some located inside or around the virus clusters (Figure 2A and Figure S3), others located outside (Table S2). Genes that are typically involved in DNA amplification (*integrase, FEN-1* and *DNApol*) and capsid formation (*p6.9, 38K* and *vp39*) in baculoviruses have been found pseudogenized in both *Campoplex* species (Table 1), except for *38K* which couldn’t be found in *C. nolae*, probably due to the higher fragmentation level of this genome. The same observation can be made in *R. megacephalus*, although, unlike in *Campoplex* and *Venturia* wasps, *p6.9* and several *integrases* have been found.

**Figure 2:**
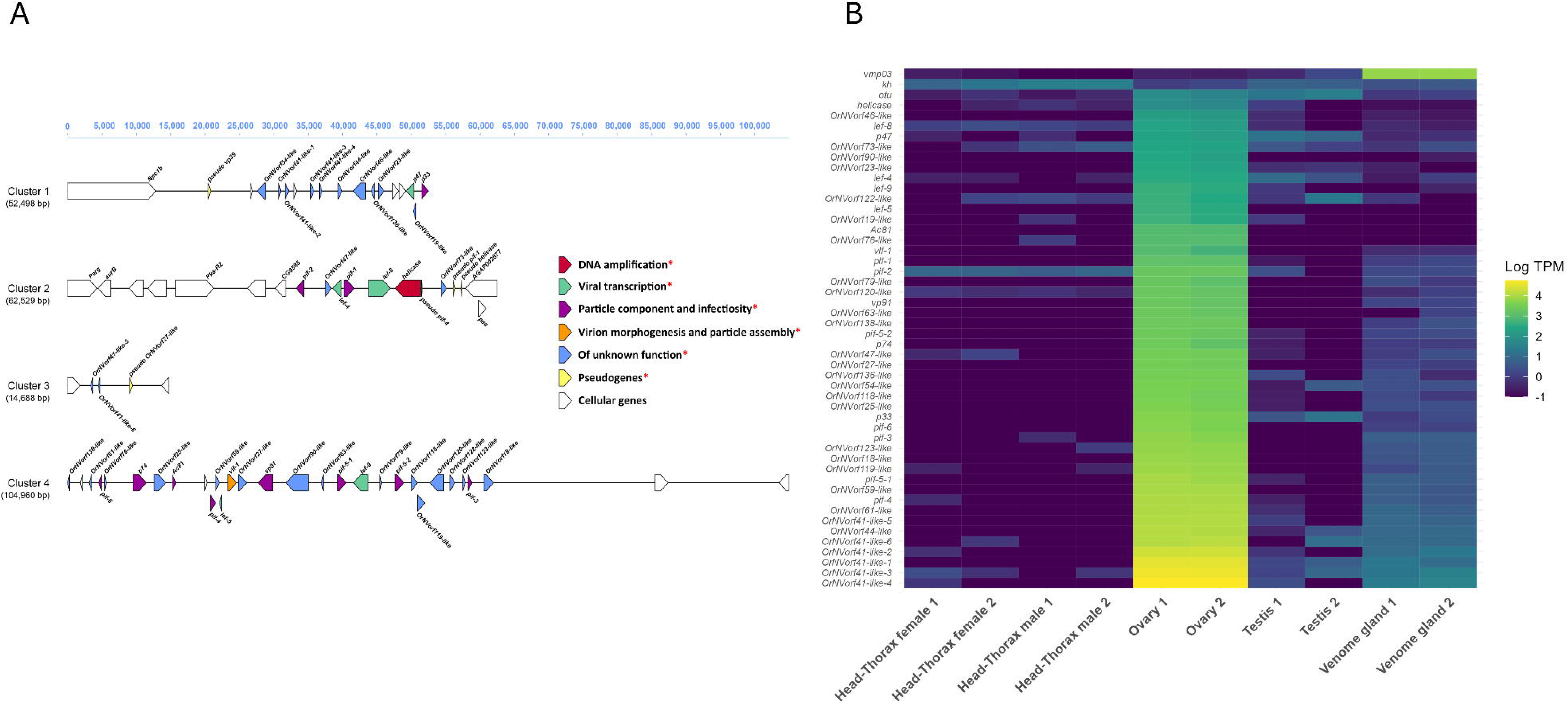
The genome of CcapiENV. A - The nudiviral clusters of CcapiENV. Sequences of nudiviral origin are indicated by a red asterisk in the legend. Nudiviral genes were colourised according to their function described in baculoviruses. Cellular genes correspond to genes of non-nudiviral origin (with no nudiviral hit). The scale represents the cluster length in base pairs. B - Expression heatmap of CcapiENV genes. Expression levels are represented by log TPMs (transcripts per million) for each organ and each replicate. The wasp genes *vmp03* (venom metalloproteinase 3), *kh* (kynurenine 3-monooxygenase) and *otu* (ovarian tumor) were added as expression controls for venom glands, head-thorax and ovaries respectively. The Spearman correlation heatmap of the transcriptome samples is available in Figure S3. An Analysis of the potential regulatory sequences of CcapiENV genes is also available in supplementary materials.

**Table 1:**
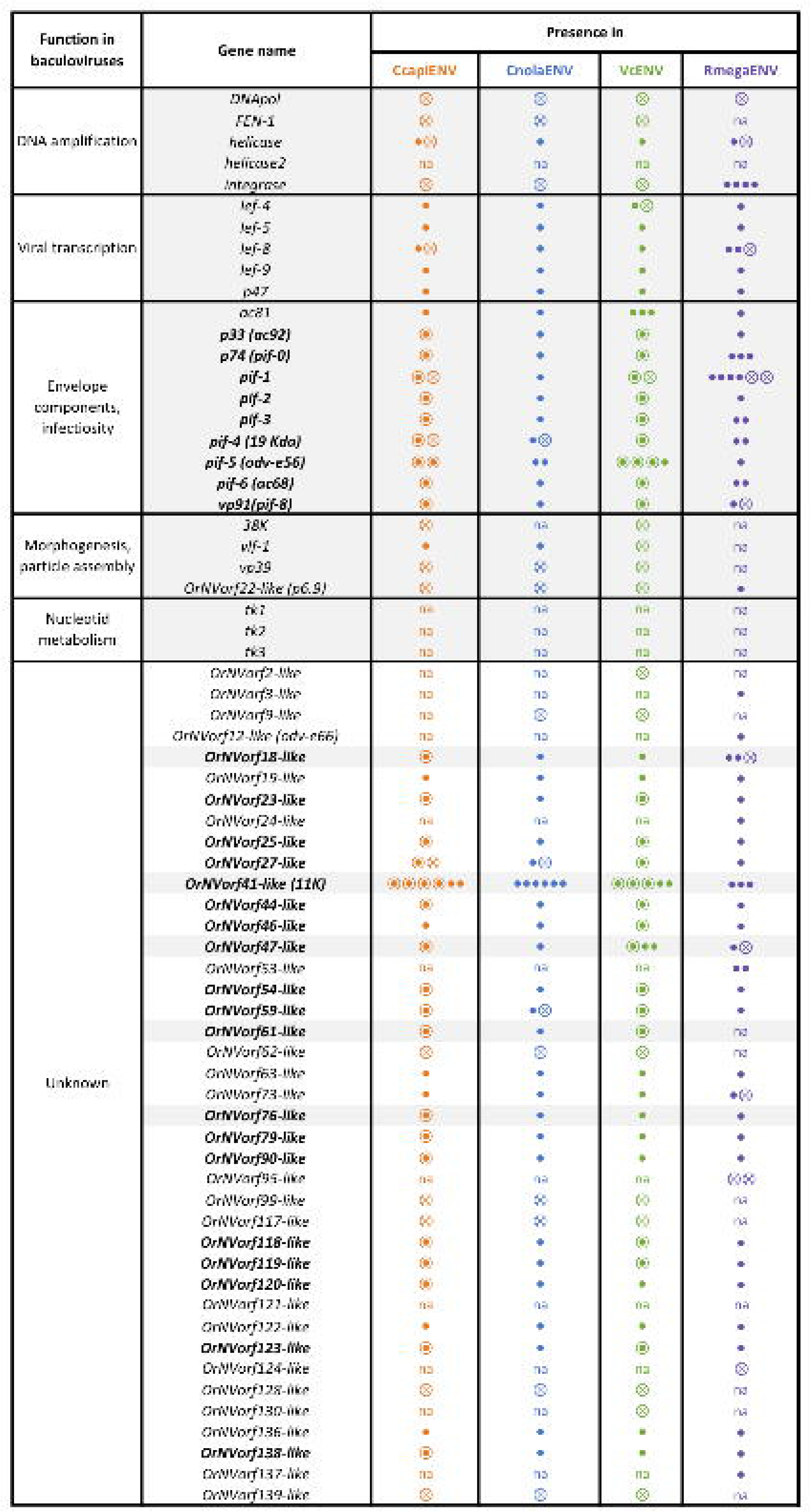
Nudivirus genes in the *C. capitator, C. nolae*, *V. canescens* and *R. megacephalus* genomes. Genes found in *C. capitator* (in orange), *C. nolae* (in blue) and *R. megacephalu*s (in purple) were compared with those of VcENV (in green). Genes and pseudogenes are indicated with solid dots and crossed dots respectively. Copies are indicated by multiple dots. Encircled solid dots indicate genes for which proteins were found in VLPs (CcapiVLPs and VcVLPs). “na” indicates that no copy of the corresponding gene was found. Alternative gene names are given between parentheses. A more detailed table is given in Table S2.

### The endogenized nudiviruses to Alphanudiviruses

Viral sequences found in *Campoplex* and *Rhimphoctona* genomes share high similarity with nudivirus sequences (Table S2). We therefore aimed to place these new endogenized viruses in the context of the phylogeny of Naldaviricetes, a class of virus comprising Baculoviridae, Nudiviridae, Hytrosaviridae and Nimaviridae. Using protein sequences from 34 genes (21 baculoviruses core genes and 13 nudivirus core genes, see Table S4 for more details), the phylogenetic analysis places these endogenized viruses inside the Nudivirus family and as a sister clade to VcENV, which belongs to the Alphanudivirus genus (Figure 3A). Due to their placement in the Nudivirus family, we called these genomes CcapiENV, CnolaENV and RmegaENV for Campoplex capitator Endogenous Nudivirus, Campoplex nolae Endogenous Nudivirus and Rhimphoctona megacephalus Endogenous Nudivirus respectively. CcapiENV and CnolaENV are from now on the closest relatives of VcENV.

**Figure 3:**
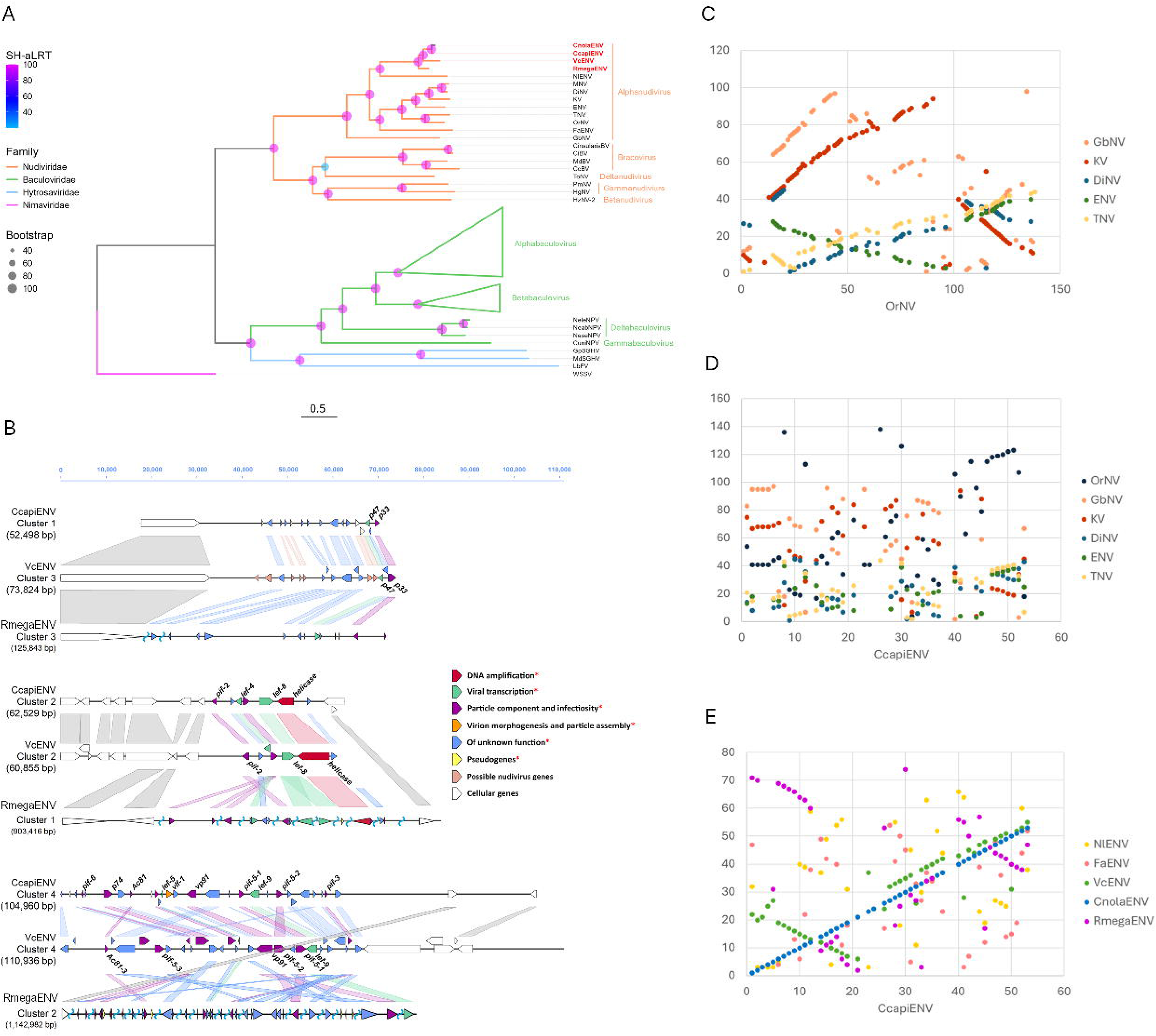
Evolution of endogenized viruses. A - Phylogenetic tree of Naldaviricetes viruses. In total, 34 genes (21 baculovirus core genes and 13 alphanudivirus core genes) were used for the tree construction, using the maximum likelihood criterion. Branch supports are given by the SH-aLRT (Shimodaira-Hasegawa approximate likelihood ratio test) and bootstrap values. The scale bar represents the phylogenetic distance (percentage of genetic variation). Campoplex capitator Endogenous Nudivirus, Campoplex nolae Endogenous Nudivirus and Rhimphoctona megacephalus Endogenous Nudivirus are called respectively CcapiENV, CnolaENV and RmegaENV. Nudiviruses endogenized in Campopleginae wasps are mentioned in red. Virus and sequence information is given in supplementary data (Table S4). B - Syntenies between the CcapiENV clusters, VcENV clusters (from Mao et al. 2023) and RmegaENV clusters. Models were colourised according to their function. Light pink models correspond to genes annotated in Mao and collaborators (Mao et al. 2023) which have no match in the NCBI database, but have nevertheless been annotated as potential virus genes. The scale represents the length in base pairs. Blue tildes on RmegaENV clusters represent genome portions that are too big to be displayed. Genome macrosyntenies can be observed in Figure S4. C - Parity plot between the genomes of Oryctes rhinoceros nudivirus (OrNV) and other exogenous nudiviruses. The other nudiviruses are Gryllus bimaculatus nudivirus (GbNV), Kallithea virus (KV) Drosophila innubia nudivirus (DiNV), Esparto virus (ENV) and Tomelloso virus (TNV). D - Parity plot between the genomes of CcapiENV and the previous exogenous nudiviruses. E - Parity plot between the genomes of CcapiENV and other endogenized nudiviruses. The other endogenized nudiviruses are Nilaparvata lugens endogenous nudivirus (NlENV), Fopius arisanus endogenous nudivirus (FaENV), VcENV, CnolaENV and RmegaENV.

### Evidence for a single nudivirus endogenization event in Campopleginae wasps

Given the close phylogenetic position of these integrated alphanudiviruses in *Campoplex* and *Rhimphoctona* (Figure 3A), we then investigated whether they resulted from the same integration event as the one for VcENV. The same wasp genes around the virus clusters were found in *C. capitator, R. megacephalus* and *V. canescens* species, indicating that these regions are homologous. This observation confirms that these viruses come from the same endogenization event that most probably occurred at the same place in a common ancestor of these wasps (Figure 3B).

### Impact of endogenization on nudivirus gene organisation

Integrated alphanudivirus genomes were compared to free virus genomes in order to evaluate the impact of viral domestication after integration. Gene order among free nudiviruses is fairly well conserved (Figure 3C), but gene order was globally lost following viral endogenization (Figure 3D), suggesting that loss of certain syntenic regions might be part of the viral domestication process.

We also compared the endogenized alphanudiviruses in Campopleginae wasps to other endogenized alphanudiviruses such as FaENV (Fopius arisanus Endogenous Nudivirus) and NlENV (Nilaparvata lugens Endogenous Nudivirus). First, as described before, the viral genome structure in Campopleginae wasps is conserved after integration (Figure 3E). However, for each independent alphanudivirus integration, the genomes seem to have been reshuffled differently (Figure 3E). Viral genome shuffling after integration might be part of a viral domestication process, as developed in Burke and collaborators, or might reflect genome rearrangements dynamic (Burke et al. 2018; Leobold et al. 2018; see discussion).

### No conclusive traces of Ichnovirus remnants in *Campoplex* genomes

As remnants of Ichnovirus were found in *V. canescens* (Pichon et al. 2015), we also searched for sequences of IVSPER genes in the *Campopex spp.* and *R. megacephalus* genomes. In *Campoplex* and *R. megacephalus* genomes, no trace of IVPSER genes could be found in using either the IVSPER protein sequences described in campoplegine wasps associated to ichnoviruses or pseudogenized IVSPER genes described in *V. canescens.* From genes potentially related to ichnoviruses, we only have detected *rep-like* genes (see Table S2), which are the most abundant genes in terms of copy number in IVs packaged genomes of campoplegine wasps. In total, while only one to two sequences have been found in *R. megacephalus* and *V. canescens*, 48 and 16 *rep-like* sequences have been found in *C. capitator* and *C. nolae* respectively. In *C. capitator*, transcripts of these sequences can be found specifically in the wasp ovaries, with some sequences showing high levels of transcription (Table S3). Given the propensity for horizontal transfer in insects (see Figure 4), we determined that the presence of *rep-like* sequences alone does not constitute evidence of ichnoviral integration.

**Figure 4:**
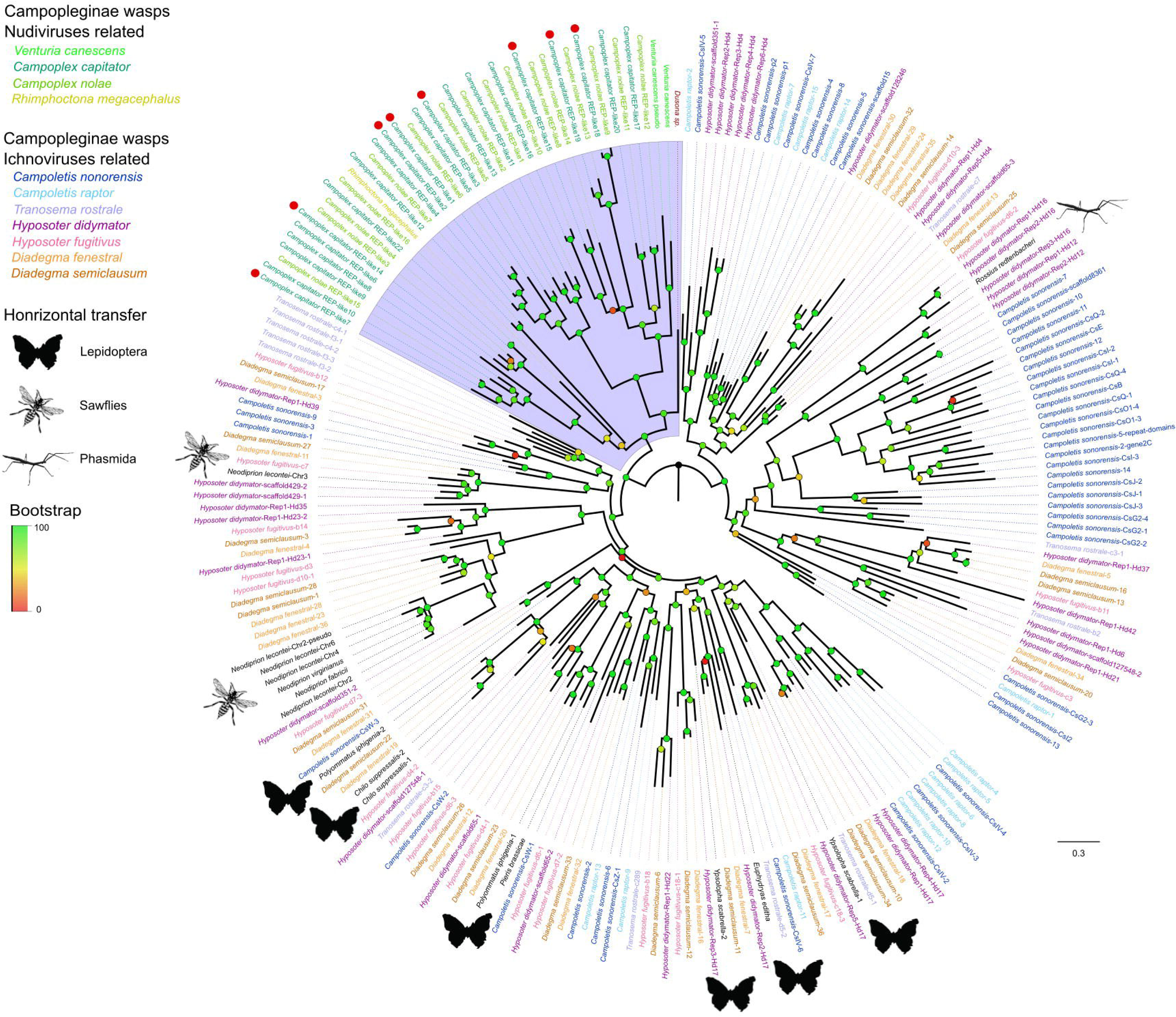
Phylogenetic tree of REP sequences across insects. The tree has been built using 229 proteins using the maximum likelihood approach. Bootstrap support values are indicated by node color. For each type of campoplegine wasp associated with an endogenous nudivirus or ichnovirus, tip label colours indicate in which species REP proteins have been identified. The pseudogenized REP identified in *Dusona sp.* is shown in red and is not associated with an active endogenous virus. Silhouettes are showing horizontal transfer of rep genes in lepidopterans, sawflies and phasmids. Red dots at the leaves indicate genes for which corresponding proteins have been found in CcapiVLPs.

### *Campoplex capitator* produces VLPs that are similar in morphology to those produced by *Venturia canescens*

After identifying the endogenized nudiviruses in the *Campoplex* genomes, we investigated whether *C. capitator* produces VLPs as described in *V. canescens*. Electron microscopy revealed the presence of particles in the calyx region of *C. capitator* ovaries, a specialised portion of the ovaries that produces virus-derived particles in braconid and Ichneumonid wasps (Bézier, Annaheim, et al. 2009; Volkoff et al. 2010; Burke and Strand 2012; Pichon et al. 2015). These particles in *C. capitator* are referred to as CcapiVLPs (Figure 5), and are almost morphologically identical to VcVLPs (Pichon et al. 2015). CcapiVLPs are composed of a membrane enclosing an electron-dense body (Figure 5E) and are massively present in the lumen (Figure 5A and B) while only a few are visible in the cytoplasm and cell nuclei (Figure 5C). Virogenic stromas, electron-dense masses composed of proteins that form viral particles in other wasps (Bézier, Herbinière, et al. 2009; Pichon et al. 2015), are visible in calyx cell nuclei (Figure 5C), along with VLPs and empty envelopes (Figure 5F). As described for VcVLPs (Pichon et al. 2015), CcapiVLPs appear to be be produced inside the nucleus by virogenic stromas and transit through the nuclear membrane and then through the cytoplasmic membrane by budding to finally end in the calyx lumen (Figure 5D,E and F).

**Figure 5:**
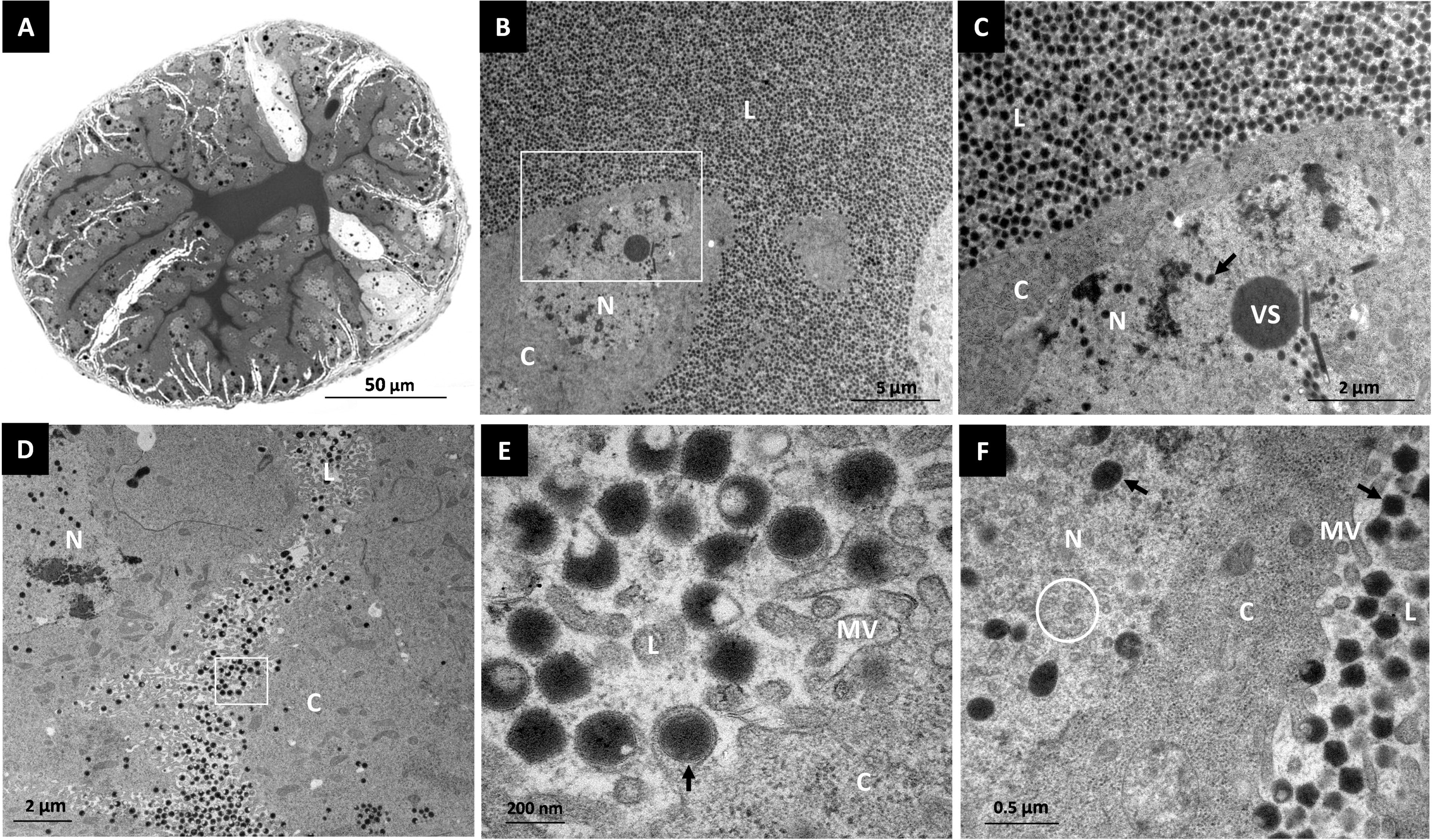
The calyx region of a *C. capitator* ovaries of adult wasp observed using light and electron microscopy. A - light microscopic photograph of a semi-thin section of the calyx region, stained with toluidine blue. B, D - electron microscopy photographs of the calyx region shown in A. C, E - enlarged areas (white frames) of photographs B and D, respectively. Virogenic stromas (VS) and VLPs (dark arrow) are visible in cell nuclei (N). While virogenic stromas can be seen only in the nuclei, VLPs appear to travel from the nucleus to the lumen (L) where they accumulate, passing through the cytoplasm (C) and the membrane microvilli (MV). F - electron microscopy photograph showing the nucleus-cytoplasm-lumen interface where empty VLP envelopes can be seen in the nucleus (white circle), as described in *V. canescens* (Pichon et al. 2015).

### *C. capitator* and *V. canescens* VLPs have a similar composition

We proceeded to determine whether the protein composition of the VLPs and their content were the same between *C. capitator* and *V. canescens*. For CcapiVLPs, 28 proteins of nudivirus origin were detected by mass spectrometry (see Table 1, Figure 6A and Table S5). In most cases, proteins contained in VcVLPs deriving from VcENV genes were also found in CcapiVLPs. For instance, all the VcVLP proteins composing the envelopes in baculoviruses were found in purified CcapiVLPs. These include all the PIF proteins, P74, VP91 and P33 proteins (Table 1, Figure 6A and Table S5). However, OrNvorf46-like and a copy of PIF-5 proteins are present in VcVLPs but not in CcapiVLPs (Table 1, Figure 6A and Table S5). Similarly, OrNVorf18-like, OrNVorf47-like, OrNVorf79-like, OrNVorf90-like and OrNVorf120-like proteins appear to be present in CcapiVLPs but were not detected in VcVLPs.

**Figure 6:**
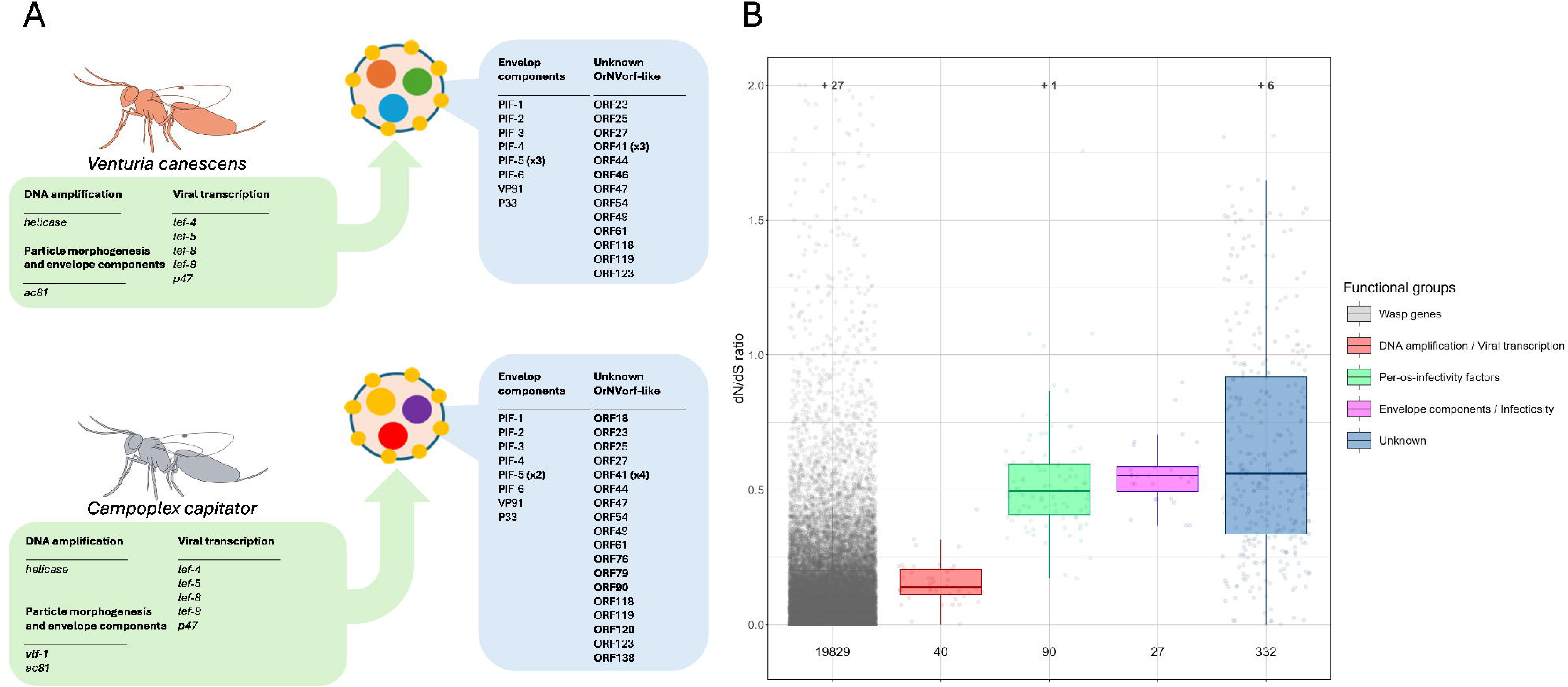
Evolution of endogenized nudivirual genes in Campopleginae. A - Protein composition of VLP found using mass spectrometry. The protein content of VLPs is given in the blue panels. In green are mentioned the nudiviral genes involved, or likely to be involved, in VLP production whose proteins are not contained in VLPs. Main differences between *V. canescens* and *C. capitator* are indicated in bold. Result details of mass spectrometry are given in Table S5. B - Estimation of selective pressures acting on wasp and endogenized nudivirus genes. dN/dS ratio values were calculated from wasp (in grey) and nudivirus genes of *C. capitator*, *C. nolae*, *V. canescens* and *R. megacephalus*. Numbers below each boxplot represent the number of sequences used for dN/dS ratio calculation for each category. Numbers above each boxplot represent the number of values higher than 2, which could not be plotted. In red are the dN/dS ratio values for *helicase*, *p47* and *lef* genes, in green are the values for *pif* (*per-os-infectivity factor*) genes, in pink the values for *Ac81*, *vp91* and *p33* genes, and in blue the values for *OrNVorf-like* genes. See Table S6 for details.

Analysis of selection pressures on *C. capitator*, *C. nolae*, *V. canescens* and *R. megacephalus* genes revealed that while 89.5% of 8,828 genes have dN/dS ratios below 1 (see Figure 6B and Table S6), nudiviral genes putatively involved in infectivity and envelope protein production have in average a higher dN/dS ratio (0.58) than genes putatively involved in DNA amplification and viral transcription (0.16). Gauthier and colleagues had also found equivalent results with nudiviral genes associated with Braconid wasps (Gauthier et al. 2018; Gauthier et al. 2021). Furthermore, duplicated nudiviral genes have on average a higher dN/dS ratio (by 1 to 2 times) than single copy nudiviral genes (Table S6).

### The virulence protein contents of VcVLPs and CcapiVLPs are different

A large number of non-viral proteins, which should include virulence proteins, could be identified by mass spectrometry within the VLPs (Table S7). Proteins described in VcVLPs were first sought in the CcapiVLP protein dataset to determine whether the same proteins are enveloped in VLPs in both wasp species. The VLP1 (PHGPx), VLP2 (RhoGAP) and VLP3 (Neprilysin) proteins of VcVLPs were not found in CcapiVLPs, indicating that VLPs produced by *C. capitator* have a different content than those produced by *V. canescens*. Candidate VLP virulence proteins were selected according to their peptide abundance (PSM for Peptide-Spectrum Matches) value and only proteins with a PSM value higher than 50 were retained (Table S8). Surprisingly, proteins encoded by eight *rep-like* genes have been found in the CcapiVLPs (Table S8). Additionally, some proteins that have been described in parasitoid wasp venoms (Poirié et al. 2014; Moreau and Asgari 2015; Inwood et al. 2023), such as a serpin and a peroxiredoxin, have been found in *C. capitator* particles. Further investigation on the serpin showed that the gene encoding this protein is a member of a multigenic family, which comprises 15 members in *Campoplex spp*. and four in *V. canescens*. While the four copies found in *V. canescens* have orthologous sequences in *Campoplex spp.*, the serpin identified in CcapiVLPs is more distant, with only one ortholog found in *C. nolae* (Figure S5). This serpin appears to be specific to *Campoplex* species and therefore represents an interesting virulence protein candidate.

## Discussion

A single, relatively recent alphanudivirus endogenization event in Campopleginae wasps

The sequencing of *C. capitator* and *C. nolae* genomes, together with analyses of *V. canescens* (Pichon et al. 2015; Mao et al. 2023) and *R. megacephalus* (Boparai and Broad 2024) genomes, has provided the opportunity to characterize the early evolutionary events following alphanudivirus domestication in Campopleginae wasps. Our results demonstrate that CcapiENV, CnolaENV, and RmegaENV derive from the same endogenization event as VcENV, as evidenced by strong synteny of flanking wasp genes around viral clusters (Figure 3B). This integration event is older than previously thought, having occurred in the common ancestor of these species. However, as Campopleginae is a younger lineage (Murphy et al. 2008; Burke et al. 2021; Sharanowski et al. 2021), this event is still more recent than the nudivirus endogenization in Microgastrinae braconids 100 million years ago (Murphy et al. 2008; Thézé et al. 2011).

Several lines of evidence support the recency of this integration. First, not all Campopleginae harbor endogenized nudiviruses; genera such as *Hyposoter* and *Dusona* either possess ichnoviruses or lack confirmed viral endogenization entirely (Burke et al. 2021), indicating that nudivirus integration occurred within a specific Campopleginae lineage rather than ancestrally. Second, viral genes in most Campopleginae wasps are organized into compact, highly conserved clusters (Figures 2A, S3, and 3B), contrasting with the more dispersed organization observed in braconid BVs (Bézier et al. 2009; Burke et al. 2014; Burke et al. 2018; Gauthier et al. 2021), suggesting fewer genomic rearrangements have occurred. Third, we identified numerous pseudogenized viral genes in *Campoplex* spp. and *V. canescens* genomes (Tables 1 and S2), whereas BV-associated wasps show complete loss of non-functional genes (Bézier et al. 2009; Burke et al. 2014). The persistence of these degraded sequences indicates insufficient time for complete erosion, consistent with previous observations in VcENV (Leobold et al. 2018).

### Genomic rearrangement as an early step in viral domestication

While gene order is highly conserved among free alphanudiviruses (Figure 3C), this synteny was largely lost following endogenization in wasps (Figure 3D). Within the same endogenization event, the endogenized viruses maintain strong synteny with each other despite lacking synteny with their free-living and endogenized relatives (Figure 3E), suggesting that major genome reshuffling occurs early in the domestication process.

This early genome shuffling may serve multiple functions in domestication. By scattering viral genes across the host genome, rearrangement could disrupt coordinated viral replication, effectively neutralizing viral pathogenicity while retaining genes useful for the wasp (Drezen et al. 2017; Burke et al. 2018; Leobold et al. 2018). However, the subsequent stabilization of these rearranged structures, as evidenced by conserved synteny across Campopleginae wasp evolution, suggests that the clustered organization may provide functional advantages for coordinated expression during VLP production. Similar conclusions have been drawn from studies of FaENV (Burke et al. 2018) and other independent endogenization events involving different virus families, indicating this may be a general feature of viral domestication.

### Conservation of essential functions and adaptive gene loss

Despite extensive genomic rearrangement, genes critical for VLP production have been selectively retained across all Campopleginae species examined. The 22 nudivirus core genes found in VcENV are present in CcapiENV and CnolaENV (Table 1), with particularly strong conservation of genes involved in viral transcription (*p47*, *lef* genes) and DNA amplification (*helicase*): these genes show significantly lower average dN/dS ratios (0.16) compared to envelope and infectivity genes (0.58) (Figure 6B, Table S6), indicating strong purifying selection. The functional importance of *lef* genes has been experimentally demonstrated, as their silencing abolishes or reduces particle production in *V. canescens* and *M. demolitor* (Burke et al. 2013; Cerqueira de Araujo et al. 2022).

In contrast, genes unnecessary for VLP production have been systematically lost or pseudogenized. Genes involved in capsid formation (*vp39*, *38K*, *p6.9*) and DNA packaging (*integrase*, *FEN-1*, *DNApol*) are pseudogenized in all VLP-producing Campopleginae wasps (Table 1), consistent with the fact that these particles lack capsids and DNA, although *R. megacephalus* retained one copy of *p6.9* and several copies of *integrase*, which could indicate that this species produces a different kind of particle. This pattern contrasts with FaENV and BVs, which retain functional copies of *38K* and *vp39* (Bézier et al. 2009; Burke et al. 2014; Burke et al. 2018), reflecting their production of DNA-containing particles.

Interestingly, *vlf-1* is functional in *Campoplex* species but pseudogenized in *V. canescens* and absent in *R. megacephalus* (Table 1). In baculoviruses, VLF-1 participates in late gene transcription and virion maturation (McLachlin and Miller 1994; Yang and Miller 1999; Vanarsdall et al. 2006). Although VLF-1 was not detected in CcapiVLPs, the gene is specifically expressed in ovaries (Figure 2B, Table S3), suggesting it may regulate the expression of some nudiviral genes involved in VLP production. Additionally, *OrNVof61-like* is absent in RmegaENV (Table 1). In *V. canescens*, this gene is lately expressed in ovaries (Cerqueira de Araujo et al. 2022) and its knockdown results in altered morphology in a subset of VLPs and increases host-mediated egg encapsulation (Mao et al. 2025). Given these differences in gene content between RmegaENV and the other endogenized viruses examined here, it is plausible that RmegaVLPs differ structurally from those produced by *V. canescens* and *Campoplex* species, which may reflect adaptation to distinct host taxa (*V. canescens* and *Campoplex* species parasitize lepidopterans, whereas *R. megacephalus* parasitizes wood-boring beetles, i.e., coleopterans). Structural divergence remains to be explored experimentally in these wasps.

### Diversification through gene duplication and divergent envelope evolution

We identified several duplicated nudivirus genes with distinct patterns across species: *pif-5* is duplicated in VcENV, CcapiENV and CnolaENV, but not in RmegaENV; *p74*, *pif-1*, *pif-3*, *pif-4*, *pif-6*, and *lef-8* are single-copy genes in *Campoplex* spp. and *V. canescens*, but duplicated in *R. megacephalus*; VcENV has its own specific duplications: *Ac81* and *OrNVorf47-like*; and *OrNVorf41-like* (an 11K family member) is duplicated in all studied species (Table 1). In baculoviruses, 11K family genes (comprising *Ac145* and *Ac150*) mediate host-specific oral infectivity (Lapointe et al. 2004; Beperet et al. 2015). Consistently, RNA interference targeting *OrNVorf41*-like-1 and -2 in *V. canescens* results in the production of seemingly empty virus-like particles (VLPs) with electron-dense membranes, and has furthermore shown that these genes are essential to escape host immunity (Mao et al. 2025). The *pif* genes encode envelope components involved in particle infectiosity, their proteins enable particle entry in the host gut cells (Kikhno et al. 2002; Zheng et al. 2017; Boogaard et al. 2018). The functions of these genes suggest that their duplication may enable wasp species to parasitize different host ranges, possibly by tailoring VLP specificity for host cell membrane proteins and/or, hypothetically, modulating the recruitment of virulence factors.

Most duplicated genes show elevated dN/dS ratios compared to single-copy genes (Table S6), indicating relaxed constraint or positive selection. This pattern is particularly pronounced for envelope and infectivity genes, which show 1- to 2-fold higher dN/dS ratios than transcription machinery genes (Figure 6B). Similar patterns have been observed in BV-associated braconids and filamentous virus-associated *Leptopilina* wasps (Di Giovanni et al. 2020; Gauthier et al. 2021), suggesting that envelope proteins evolve rapidly in response to host-specific selection pressures while core replication machinery remains conserved.

### Striking divergence in virulence protein content

Despite producing morphologically similar VLPs with nearly identical viral protein compositions (Figures 5 and 6A, Table S5), *C. capitator* and *V. canescens* package completely different virulence proteins. The three major VcVLP virulence proteins—VLP1 (PHGPx), VLP2 (RhoGAP), and VLP3 (neprilysin)—are absent from CcapiVLPs. Instead, CcapiVLPs contain a distinct set of candidate virulence factors, including a serpin, peroxiredoxin, and remarkably, proteins encoded by eight *rep*-like genes (Table S8).

The serpin identified in CcapiVLPs is particularly interesting as it belongs to an expanded gene family with 15 members in *Campoplex* species versus only four in *V. canescens* (Figure S5). This serpin is specific to *Campoplex* species and represents a prime candidate for mediating immune suppression, as serpins in other parasitoid systems inhibit the phenoloxidase cascade involved in melanisation (Beck and Strand 2007; Colinet et al. 2009). The presence of other venom-associated proteins like peroxiredoxin, which protects against oxidative stress during encapsulation (Perkin et al. 2015), could further suggest that CcapiVLPs employ fundamentally different molecular strategies for host immunosuppression.

### Convergent recruitment of rep virulence factors via distinct delivery mechanisms

The most striking discovery is the packaging of REP proteins into CcapiVLPs. *Rep* (repeat element) genes are the most abundant gene family in wasp-associated ichnoviruses, with 30 members in CsIV, 36 in HfIV, and 10 in TrIV (Galibert et al. 2006; Webb et al. 2006). While their precise function remains unclear, their conservation, high expression in parasitized hosts, and extensive expansion across IV genomes strongly suggest critical roles in parasitism success (Volkoff et al. 2002; Tanaka et al. 2007; Kim et al. 2023).

We identified 48 and 16 *rep*-like sequences in *C. capitator* and *C. nolae*, respectively, versus only one or two in *R. megacephalus* and *V. canescens* (Table S2), with several showing high ovary-specific expression (Table S3). Critically, eight REP proteins were detected in purified CcapiVLPs (Table S8), demonstrating that these virulence factors are packaged into VLPs.

This represents a remarkable example of convergent evolution: the same virulence protein family is deployed by different Campopleginae lineages using entirely different delivery mechanisms—DNA circles derived from ichnoviruses versus protein-filled VLPs derived from nudiviruses. The expansion of *rep* genes specifically in *C. capitator* reflects a similar adaptive strategy to venom protein diversification in parasitoids, whereby gene duplication enables the generation of host-specific virulence arsenals in response to ongoing selection pressures.

### No compelling evidence for virus replacement in Campoplex

Previous studies suggested *V. canescens* might have undergone “endogenous virus replacement,” losing an ancestral ichnovirus after acquiring the alphanudivirus (Pichon et al. 2015). However, we found no remnants of IVSPER genes in *Campoplex* and *Rhimphoctona* genomes, despite thorough searches using both functional IVSPER sequences from ichnovirus-associated wasps and pseudogenized IVSPERs from *V. canescens*. Similarly, Burke and collaborators (Burke et al. 2021) found no IVSPER sequences in *Dusona* sp., a close relative of *Campoplex*.

Current evidence suggests that ichnovirus integration occurred in the *Hyposoter*-*Campoletis* clade, while the “*Campoplex*-*Venturia*-*Dusona*” clade (including *R. megacephalus*) independently endogenized an alphanudivirus (present study; Burke et al. 2021; Sharanowski et al. 2021). Given that *rep*-like sequences also appear in lepidopterans and sawflies, we cannot definitively conclude that their presence alone indicates ancestral ichnovirus integration in *Campoplex* species. These sequences could originate via horizontal transfer from ichnovirus circles, as IV and BV integrations have been documented in lepidopteran hosts (Gasmi et al. 2015; Heisserer et al. 2023). Under this scenario, *rep*-like sequences in *Campoplex*, *V. canescens*, and *R. megacephalus* would reflect transfers from parasitized lepidopterans that had themselves been parasitized by ichnovirus-carrying wasps. However, no lepidopteran sequences clustered phylogenetically with these wasp REP sequences (Figure 4A), and more genome assemblies of lepidopteran hosts are needed to distinguish between horizontal transfer and ancestral ichnovirus integration.

### Implications for viral domestication and host-parasite coevolution

The comparative analysis of endogenized alphanudiviruses in Campopleginae wasps reveals key principles of viral domestication and highlights the remarkable versatility of virus-derived particle systems in parasitoid wasps. Early genomic rearrangement appears to be a hallmark of the domestication process, potentially disabling viral replication while preserving beneficial functions. Following this initial disruption, strong purifying selection maintains core particle production machinery, while envelope proteins diversify rapidly in response to host-specific pressures. Most strikingly, the virulence cargo of morphologically and structurally similar VLPs can diverge completely over evolutionary time, demonstrating extraordinary flexibility in how endogenized viruses are weaponized against different hosts. The VLP system in *C. capitator* exemplifies this versatility through its packaging of a *Campoplex*-specific serpin and expanded REP protein families—the latter representing a remarkable case of convergent evolution, as REP proteins are typically delivered via ichnovirus DNA circles in other Campopleginae lineages.

This convergent deployment of similar virulence strategies through different viral systems reflects a broader pattern across parasitoid wasps. Multiple viral families have been independently domesticated to produce particles essential for successful parasitism: deltanudiviruses generating BVs in braconids, ichnoviruses producing IVs in Banchinae and some Campopleginae, alphanudiviruses producing VLPs independently in *F. arisanus* and in the *Rhimphoctona*-*Venturia*-*Campoplex* clade, and hytrosavirus producing VLPs in *Leptopilina*. The repeated evolution of REP protein deployment—whether packaged in nudivirus-derived VLPs or delivered on ichnovirus circles—highlights how parasitoid wasps repeatedly arrive at similar molecular solutions through distinct evolutionary paths, driven by the relentless selection pressure of host immune responses.

To conclude, the genomes of four Campopleginae wasps harbour an alphanudivirus which derive from the same endogenization event in the common wasp ancestor. The description of these new endogenized nudiviruses sheds light on early evolutionary processes of viral domestication. Comparison with the closely related *V. canescens* species shows that there are mostly minute differences between the genomes of endogenized nudiviruses in Campopleginae wasps. However, major differences can be seen in the virulence protein content of VLPs which appears to be strikingly different between *C. capitator* and *V. canescens*, showing early adaptation to wasp’s hosts in the context of viral domestication.

## Material and methods

### Parasitoid model

*Campoplex capitator* wasps were obtained after emergence from the lepidopteran host *Lobesia botrana*, a vineyard pest that can grow on *Daphne gnidium* (Loni et al. 2016). Caterpillars of *L. botrana* were sampled from *D. gnidium* shrubs in the natural reserve of Migliarino-San Rossore-Massaciuccoli (Tuscany, Italy). Samplings were carried out in early July of 2019 and 2021, with wasps emerging between June and August (Scaramozzino et al. 2018). *L. botrana* nests were placed individually in test tubes and each tube was monitored every day. Emerging *C. capitator* were used for experiments.

To produce enough wasps for our experiments, a *C. capitator* rearing was maintained as described in (Lucchi et al. 2018; Benelli et al. 2020): *L. botrana* larvae were kept in plastic boxes (20cm D x 15cm W x 10cm H) with a nutritious substrate (30g of agar, 60g of sugar, 50g of alfalfa, 36g of Brewer yeast, 25g of Wesson salt, 180g of wheat sprout, 80g of casein, 4g of sorbic acid, 15g of Wanderzahnt vitamins, 2.5g of cholesterol, 2.5g of tetracyclin, 5mL of propionic acid (99,5%), 2mL of linoleic acid (95%), 5mL of olive oil and 1.5mL of distilled water). After the emergence, adults were kept in the same box for reproduction where the substrate was removed and droplets of honey were added. Once eggs were visible on the box surface, adults were removed and a nutritious substrate was added to feed the future larvae. *Campoplex capitator* females and males were meanwhile kept in plexiglass chambers (40cm D x 25cm W x 30cm H) with honey to allow their reproduction. L2 to L4 host larvae were put in the chambers for two hour parasitism periods and then individually reared into small boxes (5cm Ø x 5cm H) with nutritious substrate. These boxes were monitored every day to check for wasp emergence and honey was added when larvae were about to pupate. All insects were kept at 25°C and 45% of humidity, with a 16:8 day/night photoperiod.

Meanwhile, a female specimen of *Campoplex nolae* was collected in the wild by active sweeping at Powdermill Nature Reserve, Rector county, Pennsylvania, USA, and kept in liquid nitrogen until the time of DNA isolation.

### DNA extraction and sequencing

For *C. capitator*, DNA was extracted from one haploid male at emergence using the MagAttract® HMW DNA Kit (Qiagen). The male wasp tissues were disrupted in a 2 mL tube containing 200 µL of 1X DPBS. 20 µL of proteinase K, 4 µL of RNAse A and 150µL of buffer AL were added to the tube. The tube was then mixed carefully and incubated at 56°C at 900 rpm for 2 hours. The manufacturer’s protocol was followed from step 4 to step 16. Total DNA was finally eluted in 100 µL of TE buffer (10 mM Tris-Cl, 1 mM EDTA, pH 8.0) and quantified using the Qubit^TM^ DNA HS assay kit (Invitrogen) and the Qubit2.0 fluorometer (Invitrogen). DNA quality and contamination were checked by measurement of 260/280 and 260/230 OD ratios using the Varian Cary® 50 Scan spectrophotometer. Total DNA was then stored at 4°C. The library was prepared and sequenced using the PacBio Sequel II technology by the Gentyane platform (France).

For *C. nolae*, extraction was performed using the MagAttract® HMW DNA Kit (Qiagen), following the manufacturer’s protocol with the whole specimen immersed in extraction buffer and proteinase K overnight. An additional purification step using Agencourt AMPure XP beads (Beckman Coulter, Brea, U.S.A.) at 0.5X was used to remove small fragments. Genomic libraries were prepared for parallel sequencing in short-read (Illumina) and long-read (Nanopore) platforms: the Illumina library was prepared using the Kapa HyperPrep kit (Kapa Biosystems, Wilmington, MA). A-tailing, end-repair and ligation reactions were performed at a quarter volume relative to the standard protocol, while library amplification was performed at full volume. Custom, dual-indexing adapter-primers were used to allow for in silico de-multiplexing of each sample (Glenn et al., 2016). Following stub ligation and PCR, adapter-dimers were removed by performing a 0.8X bead cleaning using AMPure beads. The library was pooled with other samples at equimolar concentrations and sequenced at 4 nM as single lanes on Illumina NovaSeq 6000 S4 platform (2×150; Illumina Inc., San Diego, CA). The library for long-read data was prepared using the Nanopore ligation kit, complemented by reagents from the NEBNext Ultra II library prep kit (New England Biolabs, Ipswich, Massachusetts, U.S.A.), and then sequenced at a MinION platform (Oxford Nanopore Technologies, Oxford, U.K.) using the manufacturer’s specifications.

For both wasp species, raw data are available in NCBI (accession number of the BioProject: PRJNA936130).

### RNA extraction and sequencing

Emerging *C. capitator* wasps were dissected in order to obtain ovaries where CcapiVLP are produced. Venom glands and head-thorax of female adults and testis and head-thorax of male adults were also dissected to investigate expression of CcapiENV genes outside of the calyx region and in males. For all samples, two pools were analysed. Pools were composed of 5 individuals for head-thorax for both males and females, 21 female venom glands, 5 ovary pairs and 15 male testis pairs. Total RNA was extracted from the 10 pools in total using TRIzol Reagent (Invitrogen) according to the manufacturer’s instructions. Only the isopropanol step was replaced by cold absolute ethanol (3 volumes per sample) and total RNA was finally resuspended in 30µL of RNAse free water. RNA quantity of each pool was quantified using the Qubit^TM^ RNA HS assay kit (Invitrogen) designed for the Qubit2.0 fluorometer (Invitrogen). Samples of extracted RNA were then stored at −80°C.

For genome annotation, total RNA was extracted from two pools of four entire emerging female wasps using the NucleoSpin®RNA kit (Macherey-Nagel). Total RNA was finally resuspended in 40µL of RNAse free water. RNA quantity of the two pools was quantified using the Qubit^TM^ RNA HS assay kit (Invitrogen) designed for the Qubit2.0 fluorometer. Samples of extracted RNA were then stored at −80°C.

The sequencing of library preparations and the generation of paired-end reads were performed on an Illumina platform by the Novogene company (UK). Raw data are available in NCBI (accession number of the BioProject: PRJNA936130).

### Genome assembly and annotation

For *C. capitator*, genome and K-mer sizes were estimated beforehand using KmerGenie (Chikhi and Medvedev 2014). Raw reads were then corrected and trimmed using respectively Canu–correct and Canu–trim modules (Canu software v1.8) (Koren et al. 2017) in order to improve the base accuracy and to retrieve high quality sequence portions only. The genome was then assembled using the Canu–assemble module. The newly assembled genome was polished using Racon (v1.4.21) with the longest 80X raw reads to further improve the base accuracy using the longest raw reads. In parallel, *C. nolae* raw reads from Nanopore sequencing were filtered using Filtlong (v.0.2.1) before being assembled using Flye (v.2.9-b1768) (Kolmogorov et al. 2020). The produced assembly was then polished with Hypo (v.1.0.3.flye) using the Illumina reads. Scripts for *C. nolae* genome assembly are available at https://gitlab.in2p3.fr/marie.cariou/campoplex_assembly/.

The assembled genome completeness was assessed with BUSCO (Benchmarking Universal Single-Copy Orthologs, v5.4.2) (Manni et al. 2021) using Insecta and Hymenoptera lineage datasets (odb10) and genome contamination was checked with Blobtools (v1.1.1) (Laetsch and Blaxter 2017) for both genomes.

For both genomes, repeated elements were annotated *de novo* using the TEdenovo pipeline comprised in the REPET pipeline (v2.5) (Flutre et al. 2011). Two rounds of the TEannot pipeline (contained in the REPER pipeline) were then used to annotate repeated elements using existing databases. Repeated elements were then masked using Bedtools (v.2.29.1) (Quinlan and Hall 2010) before gene prediction. Gene models were generated for *C. capitator* using BRAKER-2 (v2.1.6) (Brůna et al. 2021) with assembled transcriptomes by spades (v3.15.5) (Bushmanova et al. 2019). Models were then functionally annotated using InterProScan (v5.53-87.0) (Blum et al. 2021) and BLASTp (NCBIblast+ v2.13.0) with the nr database (downloaded the 14^th^ of April 2023), before being polished by AGAT (v1.0.0) (Dainat et al. 2023). Proteins were extracted from gene models using the agat_sp_extract_sequences.pl tool from AGAT software. Finally, the genome completeness was assessed again using the protein sequences with BUSCO.

In total, 44 CcapiENV genes were annotated with the automated annotation process. Using the gene models predicted previously, additional viral genes were then annotated by reciprocal blasts: translated sequences from predicted genes were aligned using BLAST tools on the NCBI nr database filtered for sequences of endogenous and exogenous nudiviruses. Aligned sequences were then extracted and realigned on the complete NCBI nr database. The final BLAST outputs were filtered to keep only sequences which aligned on sequences of endogenous and exogenous nudiviruses reported in the database.

To annotate nudivirus genes from *C. nolae* genome, open reading frames were predicted using orfipy (v0.0.4) (Singh and Wurtele 2021). Nudivirus genes were then annotated by reciprocal blasts using translated sequences from predicted ORFs. Miniprot (v0.9-0) (Li 2023) was also used to align *C. capitator* proteins on *C. nolae* genome and validate previously annotated nudivirus genes. Miniprot gene models were also used for gene clustering in the dN/dS ratio analysis.

Genome assembly and gene predictions for *R. megacephalus* were obtained from the DToL data portal (accession number: GCA_963989365.1,). To annotate nudivirus genes in the *R. megacephalus* genome, a complete list of nudiviral proteins was used. Nudiviral proteins were searched against the wasp genome using tBLASTn (matrix: BLOSUM62; maximum E-value: 1e-2). All hits were then analyzed individually, and open reading frames were predicted using Geneious Prime® 2019.0.4.

Pseudogenized genes were identified by reciprocal blasts: tBLASTn was used first to map nudivirus proteins on the wasp genomes. Hits were then verified by BLASTx to map potential nudiviral nucleotide sequences on the NCBI nr database. Proteins corresponding to nucleotide sequences mapping back on nudivirus were kept and incomplete and non-transcribed sequences were considered as pseudogenes.

*Rep* genes and ichnovirus sequences were searched in *C. capitator*, *C. nolae*, *V. canescens*, *R. megacephalus* and *Dusona sp.* Genomes. Sequences from ichnoviruses were first downloaded from NCBI databases and manually completed. IVSPER sequences were searched using a homemade pipeline: genomic IVSPER sequences were first aligned on the genomes (the genome sequences were segmented into blocks of 500,000 bases) using BLASTn and tBLASTx. Aligning chunks were then aligned with BLASTx on the NCBI nr database. These hits were filtered to keep only hits aligning on IVSPER proteins. These hits were then verified by aligning them back on the nr database using BLASTp. In parallel, pseudogene sequences of IVSPER described in *V. canescens* were aligned on the genomes using tBLASTn. Protein sequences of the matching regions were then extracted and blasted against a portion of the NCBI nr database specific to Ichneumonidae (taxid:7408) and Viruses (taxid:10239) using a PSI-BLAST with an e-value threshold and a word size set to 0.5 and 2 respectively. Finally, only hits aligning with IVSPER genes contained in the NCBI database were retained. *Rep* genes and pseudogenes were all annotated manually using tBLASTn.

### CO1 and nudivirus gene alignments

Because the endogenized nudivirus genomes were highly similar in both *Campoplex* species, we collected evidence to confirm that the *Campoplex* wasps sequenced for this study are part of different species. Methods and results for this part are available in the supplementary materials.

### Campopleginae and endogenized nudivirus evolution

To obtain a phylogenetic tree of ichneumonid wasps, complete BUSCO sequences from RefSeq assemblies were extracted using BUSCO and the hymenoptera odb10 database. Sequences were then aligned using MAFFT (v7.487) (Katoh et al. 2019) and the following parameters: maxiterate=1000 (maximum number of iterations), genafpair (Altschul algorithm (Altschul 1998)). Aligned sequences were then trimmed by TrimAL (v1.4.rev15) (Capella-Gutierrez et al. 2009) with the gap threshold parameter fixed at 0.6. Trees were then generated for each sequence using IQ-TREE 2 (v2.1.4) (Minh et al. 2020) and the following parameters: m=TEST (use the best model found), B=1000 (number of bootstrap iterations), alrt=1000 (number of SH-aLRT replicates). A species topology was then inferred using ASTRAL (v 5.7.8) (Zhang et al. 2018). To build the final species phylogeny, sequences were concatenated using goalign (v 0.3.7) (Lemoine and Gascuel 2021) and a partitioned tree was built using IQ-TREE 2 with the constrained topology inferred by ASTRAL.

To perform the virus phylogeny, protein sequences of one nimavirus, three hytrosaviruses, sixty-nine baculoviruses and twenty nudiviruses (including CcapiENV and *C. nolae* sequences) were first aligned together to find orthologous genes. Results were then manually corrected using the knowledge found in the literature (Bézier, Annaheim, et al. 2009; Bézier, Herbinière, et al. 2009; Burand et al. 2012; Bézier et al. 2015; Burke 2019; Drezen et al. 2022). Protein sequences of orthologous genes were then aligned using MAFFT with the previous parameters. Alignments were then trimmed by TrimAL (gap threshold = 0.6). For each gene, a tree was finally generated with IQ-TREE 2 with the previous parameters. Protein sequences of genes that were not supporting the accepted phylogeny (at the family level) were removed from the further analysis. Finally, 21 baculovirus core genes and 13 alphanudivirus core genes were kept for the virus tree construction. Protein sequences of these genes were aligned and trimmed as described previously. Sequences were then concatenated using goalign and a partitioned tree was constructed with iqtree2 with the same parameters as previously described. The tree image was then built using ape (v5.7-1), treeio (v1.24.0) (Wang et al. 2020) and ggtree (v3.8.0) (Yu et al. 2017) packages with R (v4.2.1).

A whole genome synteny analysis was performed using the R package GENESPACE (v1.2.3) (Lovell et al. 2022). Parity plots (Figure 3C to E) showing syntenies among nudiviruses were generated with Excel (Microsoft v16).

To estimate the evolutionary constraint occurring on groups of genes according to their function, protein sequences of *C. capitator*, *C.nolae*, *V. canescens* and *R. megacephalus* were first clustered using OrthoMCL (v2.0.9) (Li et al. 2003) and scripts written by Arun Seetharam (https://github.com/ISUgenomics/common_scripts). Clustered protein sequences were then aligned using ClustalOmega (v1.2.4) and aligned sequences were converted into codon alignment using PAL2NAL (v14.1) (Suyama et al. 2006). Phylip trees were generated using IQ-TREE 2 with the following parameters: -m JTT (Jones-Taylor-Thornton model (Jones et al. 1992)). Codeml (Paml v4.10.6) was finally used to calculate dN and dS values.

To construct the phylogenetic tree of *rep* genes, *rep* sequences from ichnovirus-associated wasps and newly annotated *rep* were analysed altogether with *rep* sequences identified in lepidopterans, sawflies, and phasmids - which were all annotated manually using tBLASTn. Amino acid sequences were then aligned with MAFFT using parameters adapted for a divergent gene family (--localpair, --maxiterate 1000, --bl 45). The phylogenetic tree was subsequently generated using IQ-TREE 2 (v2.4.0) with the following parameters: -m MFP, -bb 1000, -alrt 1000. Sequences forming long branches and a group of REPs in *C. capitator* were removed, and the entire analysis was repeated on the remaining sequences. The sequences forming long branches were excluded to avoid artifacts in the tree topology. The group of REPs in *C. capitator* corresponded to a set of sequences showing 100% similarity among small genomic scaffolds, which were excluded due to a suspected assembly bias. To simplify the final figure, only bootstrap values are shown (Figure 4).

### Gene expression and regulatory sequences of CcapiENV genes

Results and methods are available in the supplementary materials.

### CcapiVLP purification and protein sequencing

In order to purify Virus-Like-Particles, 61 adult female wasps of *C. capitator* were dissected 2 days after emergence. In total, 122 calyces were obtained and pooled in 1X DPBS. Calyces were then disrupted thanks to a pellet pestle and CcapiVLPs were purified as described in (Pichon et al. 2015): disrupted calyces were centrifuged at 770 g, at 4°C for 10 minutes in order to separate CcapiVLPs and calyx fluid (supernatant) from cellular waste (pellet). The supernatant was then retrieved and transferred into a new tube. The new tube was then centrifuged at 15 400 g, at 4°C for 10 minutes. The supernatant was discarded and the pellet containing CcapiVLPs was suspended in 20 µL of 1X DPBS and stored at −20°C.

CcapiVLP proteins were then separated by SDS-polyacrylamide gel electrophoresis (12.5% polyacrylamide gel) under denaturing conditions. Separated bands were incubated for 1h with a Coomassie blue solution (10% acetic acid, 50% ethanol, 0.1% R250 Coomassie blue, q.s. to 1L of milliQ water) and 9 slices were cut from the gel. Slices were chopped into 1mm^3^ cubes before being stored at −20°C and analysed by the platform PIXANIM (France) using a nanoUPLC_LTQ-VElos Pro Orbitrap Mass Spectrometer.

For each protein identified by the mass spectrometry, sites and domains were searched for using InterProScan and gene ontology terms were attributed using sequence homology with curated proteins from the Uniprot SWISSprot database (downloaded the 18th November 2022). Proteins with a peptide abundance higher than 50 were then retained as VLP candidates.

### Electronic microscopy of the calyx region

The calyx region of adult *C. capitator* wasp ovaries was observed under electronic microscopy. Wasp ovaries were dissected two days after emergence and fixed in PrOx/EPON epoxy resin. Samples were fixed, cut and observed with the same protocol as one used in Cerqueira de Araujo and collaborators (Cerqueira de Araujo, Leobold, et al. 2022). The protocol is available in Supplementary materials.

## Acknowledgements

The authors would like to thank Francesca Cosci, Filippo Di Giovanni and Augusto Loni from the Department of Agriculture, Food and Environment (DAFE) of the University of Pisa (Italy) for the help provided during field wasp sampling. The authors would like to thank also Amine Chebbi who helped the first author on the transposable element annotation pipeline and Christophe Bressac who determined the sperm production period of male *C. capitator*. We would like to thank the IBiSA Electron Microscopy Facility of the University of Tours and the University Hospital of Tours for their assistance. Research funding was provided by an ADS Research Grant from the Smithsonian Institution to BFS, who was also supported by a GGI Peter Buck postdoctoral fellowship during early stages of this work.

## Data availability

Raw reads and assemblies are deposited on the NCBI database under BioProject PRJNA936130.

