## Supplementary Materials for "Rep virulence genes have been recruited convergently by nudivirus-derived Virus-Like Particles and ichnoviruses of Campoplegine wasps"

##### Species determination

To determine if the genomes of *C. capitator* and *C. nolae* come from distinct species, whole genome alignments were performed using D-Genies (v1.4.0) (Cabanettes and Klopp 2018). A CO1 (cytochrome oxydase 1) sequence alignment was also performed in order to measure divergence between the two genomes. CO1 orthologs were searched for in *C. capitator* and *C. nolae* genomes by aligning ORFs on CO1 sequence of *Campoplex* species reported in BOLD database (Ratnasingham and Hebert 2007). Candidates were then aligned together (*C. capitator* candidates vs *C. nolae* candidates) before being aligned on the NCBI database. Endogenized nudivirus were also aligned by blastp to assess the protein divergence between the two species.

As the endogenized nudivirus genomes were highly similar in terms of structure and sequence identity, we gathered pieces of evidence confirming that the two *Campoplex* specimens sequenced herein come from two distinct wasp species. To determine whether the two species belong to different species of *Campoplex*, genome comparison was performed between the two species. At the genome scale, the identity stands predominantly between 50% and 75% (Figure S4). 11% of these genomes share an identity higher than 75%. In comparison, using D-Genies, we found that *Cotesia flavipes* and *Cotesia sesamiae*, two closely related species of braconid wasps, share 57% of their genome with an identity standing between 50% and 75%, and 21% with an identity higher than 75%. This indicates that two different species can share high levels of similarities and remain two distinct species.

CO1 gene identification revealed that the protein sequence from the two species shares 90.68% of identity with 48 mismatches and, while *C. nolae* CO1 matches at 100% of identity with a *Campopleginae sp.* sequences (GenBank MG343679.1), *C. capitator* CO1 matches at 100% of identity with a sequence of *Campoplex difformis* (GenBank: MZ622859.1). It should be noted that *C. capitator* have been also called *C. difformis* due to identification issues (Scaramozzino et al. 2018).

The pairwise identity between CnolaENV, CcapiENV and VcENV amino acid sequences was also used to approximate the divergence level between the genomes. The average pairwise identity between CnolaENV and CcapiENV is 93%. The amino acid sequence of the *pif-1* gene harbours the lowest identity between the two nudiviruses endogenized in *Campoplex* (82.7%), while the one of *lef-8* possesses the highest identity (98.1%). If we look at braconid wasps harbouring a bracovirus, blast alignment of LEF-8 protein sequences gives an identity ranging between 70% and 99%. The LEF-8 sequence of *C. glomerata* shares 98.95% identity of the one of *C. congregata*. On the other hand, PIF- 1 shares between 30% and 77% identity in braconids. The PIF-1 protein of *C. glomerata* shares with the one of *C. congregata* 76.95% identity.

All these clues led us to the conclusion that *C. capitator* and *C. nolae* are most probably not part of the same species, although they are closely related.

Partial sequences of CO1 gene of *C. capitator* and *C. nolae*:

>*Campoplex capitator* CO1 (partial)

ATGGGATCTCCTCCTCCAGAAGGATCAAAAAAAGAAGTATTTAAATTACGGTCTGTTAATAATATAGTGATTGCTCCTGCTAAAACAGGAACAGCTAGTAATAATAAAATAGTTGTAATTAAAATTGATCATGTGAATAAAGTTAAATTTTCTAAATTTTTATTAATATTTTTTATTAAAAAAATAGTTGAAATAAAATTAATTGCTCCTATAATAGAGGATATTCCTGCGATATGTAATGAAAAAATAGCTAAATCTACTGATATACCTTCATGATTAATATTTAAAGATAAAGGAGGATATACTGTTCATCCTGTTCCAGCTCCTTGATTAATAATGGATCTTGATATAAGTAATAAGATAGAAGGAGGTAATAATCAAAATCTTATATTATTTATACGGGGAAATGCTATATCAGGTCTTCCTATTATTAGTGGAATTAATCAATTTCCAAATCCTCCAATTATAATTGGTATAACTATAAAGAAAATTATAATAAATGCATGTGCAGTTACAATTGAATTATAA

>*Campoplex nolae* CO1 (partial)

ATAGGGTCTCCTCCTCCAGATGGGTCAAAAAAAGAAGTATTTAAATTTCGATCTGTTAATAATATAGTAATTGCACCAGCTAAAACTGGAACAGCTAATAGTAATAGAATAGTTGTAATTATGATTGATCATGTAAATAGTGTTATTTGTTCTAAATTTTTATTAATATTTTTTATTAATAAAATAGTTGAAATAAAATTAATTGCTCCTATAATTGATGATATACCTGCAATATGTAAAGAAAAAATGGCTAGATCTACAGATATTCCTTCATGATTAATGTTTAAAGATAAAGGGGGATATACTGTTCATCCTGTTCCTACTCCTTGATTAATAATTGATCTTGATATAAGTAATAAAATAGAAGGGGGTAGTAATCAAAATCTTATATTATTTATTCGAGGGAAGGCTATGTCAGGACTTCCTATTATTAATGGAATTAATCAATTTCCAAACCCTCCAATTATAATTGGTATAACTATAAAAAAAATTATAATAAATGCATGGGCAGTTACAATTGAGTTATAA

##### *C. capitator* transcriptomes and CcapiENV gene expression

Virus gene expression activity within the wasps has been investigated through transcriptome analyses performed on *C. capitator* samples of pools of organs from male and female individuals. Paired raw reads from ovaries, head-thorax, testes and venom gland of *C. capitator* wasps were first trimmed using Trimmomatic (v 0.38) (Bolger et al. 2014) and TruSeq3-PE adaptors. A quality check was then performed on trimmed reads using fastqc (v 0.11.9) before mapping. Reads were mapped on the genome using Hisat-2 (v2.2.1-3n) (Kim et al. 2019) after genome indexation. Mapped reads were then sorted using samtools (v1.16). Sorted reads were converted into counts by FeatureCounts from the Rsubread package (v2.14.1) (Liao et al. 2019). Using the edgeR package (v 3.42.2) (Robinson et al. 2010), genes with a count being equal to less than 15 per sample were filtered from the dataset in order to remove non-expressed genes. The dataset was then normalised by converting the read counts into TPM (transcripts per million). A Spearman correlation heatmap was performed to verify duplicate consistency (Figure S2).

Because virus genes participate in the production of virus-derived particles in the ovaries for several wasp species, notably in *V.canescens* (Stoltz 1990; Drezen et al. 2006; Pichon et al. 2015), we assumed that CcapiENV genes would be mostly expressed in *C. capitator* ovaries. As expected, all CcapiENV genes are expressed in adult wasp ovaries (Figure 2B and Table S3). Transcripts from some nudivirus genes are also visible in venom glands. The head-thorax samples of female wasps appear to be almost free of viral transcripts, with only minute amounts of transcripts being measured. CcapiENV genes are practically not expressed in males, neither in testis nor in head-thorax samples, although testis samples show a slightly higher number of transcripts than head-thorax samples (Table S3). The *pif-2* gene appears to be an exception as it is weakly expressed in head-thorax samples of both males and female wasps.

##### Regulatory sequences in CcapiENV

In the same fashion as described in (Cerqueira de Araujo et al. 2022), promoter sequences regulating the early and late expression of viral genes in baculoviruses (Passarelli and Guarino 2007; Chen et al. 2013; Rohrmann 2014) were sought in the 300 bp upstream of CcapiENV genes. The “early” promoters described in Baculoviruses include a TATA box (TATA[A/T]T[A/T]) and a sequence motif (CA[G/T]T or CGTCG) placed among the 40 nucleotides after the TATA box. The late promoters include the baculovirus motif ([A/T/G]TAAG) and the Heliothis zea nudivirus 1 motif (TTATAGTAT) (Cheng et al. 2002). We also considered the “late” motif ([G/T][A/T][A/G]A[A/T]ATAG[T/A]) described in Alexandra Cerqueira 2022 for which 2 mismatches were allowed in the sequence search.

Between CcapiENV and VcENV hypothetical regulatory sequences, no congruent results could be observed as the type of promoter (baculovirus early, baculovirus late, VcENV late) does not necessarily match the expected expression kinetics of the CcapiENV genes based on the transcription kinetic described for their VcENV orthologs (Cerqueira de Araujo et al. 2022); Table S9).

##### Electronic microscopy

Sample fixation occurred in a mixture of 2% paraformaldehyde (Merck, Darmstadt, Germany), 2% glutaraldehyde (Agar Scientific, France) and 0.1 sucrose in 0.1M cacodylate buffer (pH 7.4). Samples were incubated for 24 hours, before to be washed three times for 30 minutes with a washing solution (0.1M of cacodylate buffer) and post-fixed for 1.5 hours with 2% osmium tetroxide (Electron Microscopy Science, USA) in 0.1M cacodylate buffer. After being washed with the washing solution for 20 minutes plus two times for 20 minutes in distillated H_2_O, samples were dehydrated in a graded series of ethanol solutions (50% ethanol two times for 15 minutes; 70% ethanol two times for 15 minutes and third portion of 70% ethanol for 14 hours; 90% ethanol two times for 20 minutes; and 100% ethanol three times for 20 minutes). Final dehydration was performed by 100% propylene oxide (PrOx, VWR Int., France) three times for 20 minutes. Samples were then incubated in PrOx/EPON epoxy resin (Fluka, Switzerland) mixture in a 2:1 ratio for two hours with closed caps, in PrOx/EPON epoxy resin (Fluka, Switzerland) mixture in a 1:2 ratio for two hours with closed caps and 1.5 hours with open caps, and in 100% EPON for 16 hours at room temperature. Samples were replaced in new 100% EPON and incubated at 37 °C for 24 hours and at 60 °C for 48 hours for polymerization.

Semi-thin sections (thickness 0.8 µm) of ovaries (cross sections) were cut with a “Leica Ultracut UCT” ultramicrotome (Leica Microsysteme GmbH, Wien, Austria), placed on glass, stained with Toluidine blue (Electron Microscopy Science, Hatfield, PA, USA) and embedded in Epon resin (Fluka, Switzerland) which was allowed to polymerize for 48 hours at 60°C. The sections were then observed with Nikon Eclipse 80i microscope (Nikon, Japan) connected to DS-Vi1 camera driven by Nis-Element D 4.4 imaging software (Nikon, Japan).

Ultra-thin sections (thickness 70 nm) were cut with a “Leica Ultracut UCT” ultramicrotome (Leica Microsysteme GmbH, Wien, Austria), placed on TEM one-slot grids (Agar Scientific, France) coated with Formvar film and stained 20 minutes with 5% uranyl acetate (Electron Microscopy Science, Hatfield, PA, USA) and 5 minutes Reynolds lead citrate. The sections were then observed at 100 kV with a Jeol 1011 TEM (Tokyo, Japan) connected to a Gatan digital camera driven by Digital Micrograph software (GMS 3, Gatan, Pleasanton, CA, USA).

### Figures and tables

Table S1: *Campoplex* genome metrics. A - Genome metrics after assembly and polishing. Metrics were obtained by Quast. B - *Campoplex* genome completeness assessed by BUSCO. “Insecta” and “Hymenoptera” refer to the OrthoDB 10 used to perform the analysis. Genome completeness was assessed at the genomic scale (--mode option = “genome”) and the gene content scale (--mode option = “protein”) after gene model prediction.

Table S2: Summary table of CcapiENV (*C. capitator*), CnolaENV (*C. nolae*), RmegaENV (*R. megacephalus*) and *rep* genes information. Nudivirus and *rep* genes were validated by blastp. Nudivirus core genes are indicated in bold. REP (repeat element proteins) ichnovirus functions are underlined. Alternative gene names are given between parentheses. TPMs (transcripts per million) were generated from ovary transcriptomes and the presence of a given protein in VLPs was determined by mass spectrometry. PSM corresponds to peptide-spectrum matches. An “*” indicates that the accession mentions the protein between parentheses according to NCBI GenPept. the “**” corresponds to the same amino acid sequence as AJZ73152.1, which is *Venturia canescens* OrNVorf138-like.

Table S3: TPMs of CcapiENV and *rep* genes per organ. “HT”, “ov” and “vg” correspond to head-thorax, ovary and venom gland samples respectively. Each sample corresponds to a pool of several individuals.

Table S4: Accession numbers of sequences used for the tree generation with IQ-TREE.

Table S5: Viral protein sequences contained in purified VLPs identified by mass spectrometry. For each protein, peptide sequences identified by mass spectrometry are coloured in red. Peptide abundance is measured in PSM (peptide-spectrum matches). Results obtained from purified CcapiVLPs were compared to results obtained from purified VcVLPs (Pichon et al. 2015).

Table S6: dN/dS ratios of *C. capitator*, *C. nolae, R. megacephalus* and *V. canescens* genes.

Table S7: Proteome of the CcapiVLP sample identified by mass spectrometry. Only proteins of cellular origin (non-nudiviral) were reported in this dataset.

Table S8: List of the most represented proteins in CcapiVLPs. Proteins were sorted according to their peptide abundance (PSM) and only proteins with a PSM superior to 50 were kept. REPs and proteins sharing function similarities with parasitoid wasp venom proteins were highlighted in light blue (see Poirié et al. 2014; Moreau and Asgari 2015; Inwood et al. 2023 for references). The full version of this table is given in Table S9.

Table S9: Possible regulatory sequences identified in the upstream region of CcapiENV genes. ^a^corresponds to the expression kinetic of VcENV described in (Cerqueira de Araujo et al. 2022). Dots correspond to regulatory sequences of early and late expressed genes described in baculoviruses (filled dot and encircled dot), to a motif found in HzNV-1 (split dot) and to a motif found upstream late expressed genes in VcENV (hashed dot). Results were compared to regulatory sequences found in VcENV.

Figure S1: Phylogenetic tree of ichneumonid wasps. The tree was built using hymenopteran BUSCOs extracted from NCBI genome assemblies. Branch topology has been inferred by coalescent modelling (ASTRAL) while branch lengths were inferred using a maximum likelihood approach. Branch support is given by local posterior probabilities, i.e. the probability that a branch is the true branch given the set of trees built using BUSCOs (computed based on the quartet score (Sayyari and Mirarab 2016)). Wasps having integrated viruses in their genome are highlighted in blue.

Figure S2: Spearman correlation heatmap built from transcriptome samples. “HT”, “ov” and “vg” correspond to head-thorax, ovary and venom gland samples respectively. For each condition, replicates cluster together, the intra-condition variation appears to be higher than the inter-condition variation.

Figure S3: The nudiviral clusters of CnolaENV and RmegaENV. Genes were colourised according to their function described in baculoviruses. The scale represents the cluster length in base pairs. Portions of RmegaENV clusters were zoomed (highlighted in blue) for better readability (skipped sequences are indicated with a blue tilde).

Figure S4: Synteny analysis between *C. capitator*, *C. nolae*, *V. canescens* and *R. megacephalus* genomes. A/ Syntenies between all genomes. Colours match *V. canescens* chromosomes. B/ Positions of shared nudivirus clusters across all genomes. Figures were generated using the R package GENESPACE.

Figure S5: Phylogenetic tree of the serpin family including the serpin found in CcapiVLPs. Serpin protein sequences were used to generate the tree. The serpin found in CcapiVLPs is indicated in red. Values correspond to bootstrap support values.
