## Supplementary figures and images for "Rep virulence genes have been recruited convergently by nudivirus-derived Virus-Like Particles and ichnoviruses of Campoplegine wasps"

### Figure S1

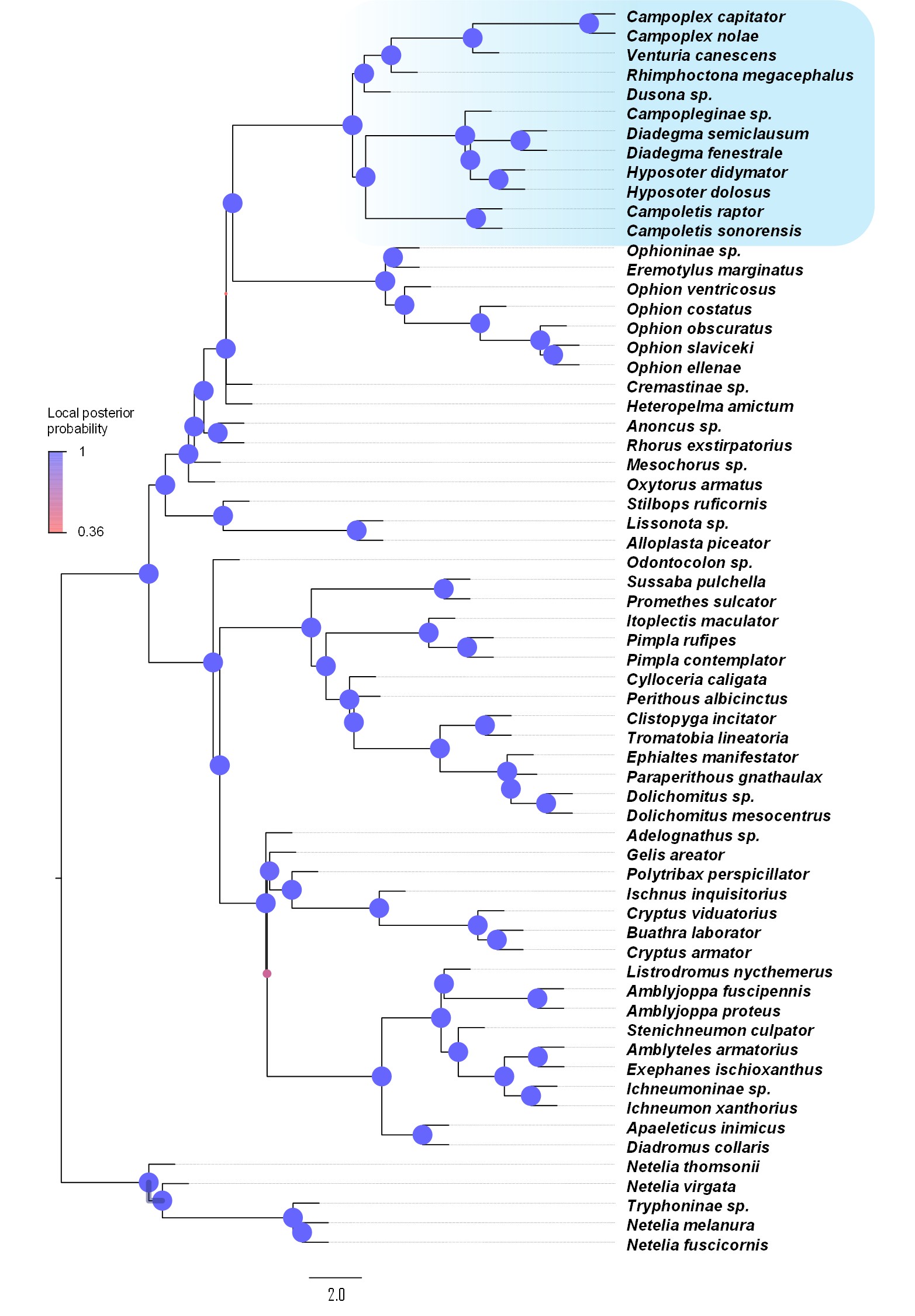

### Figure S2

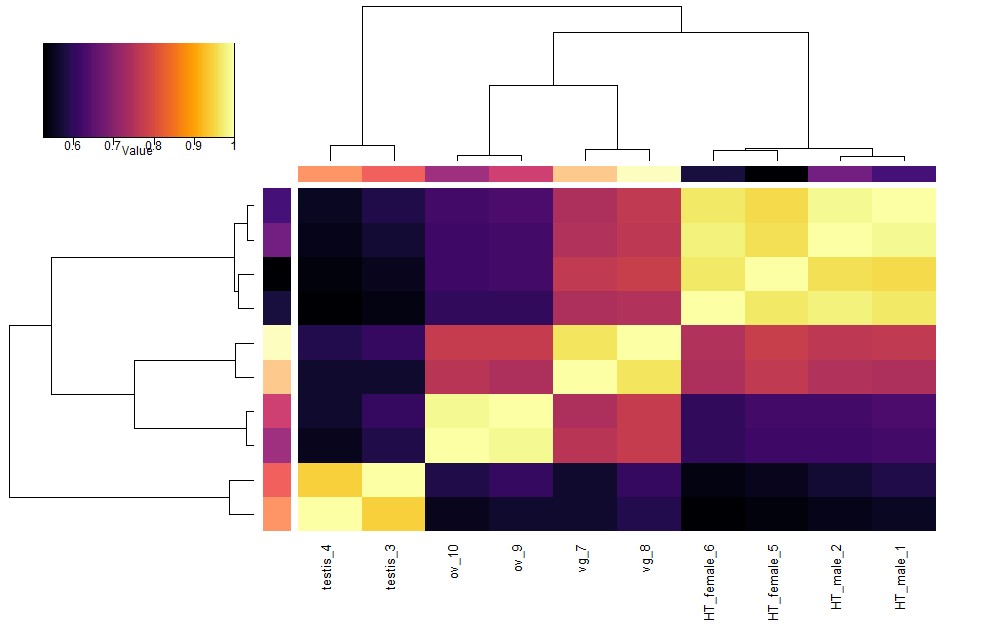

### Figure S3

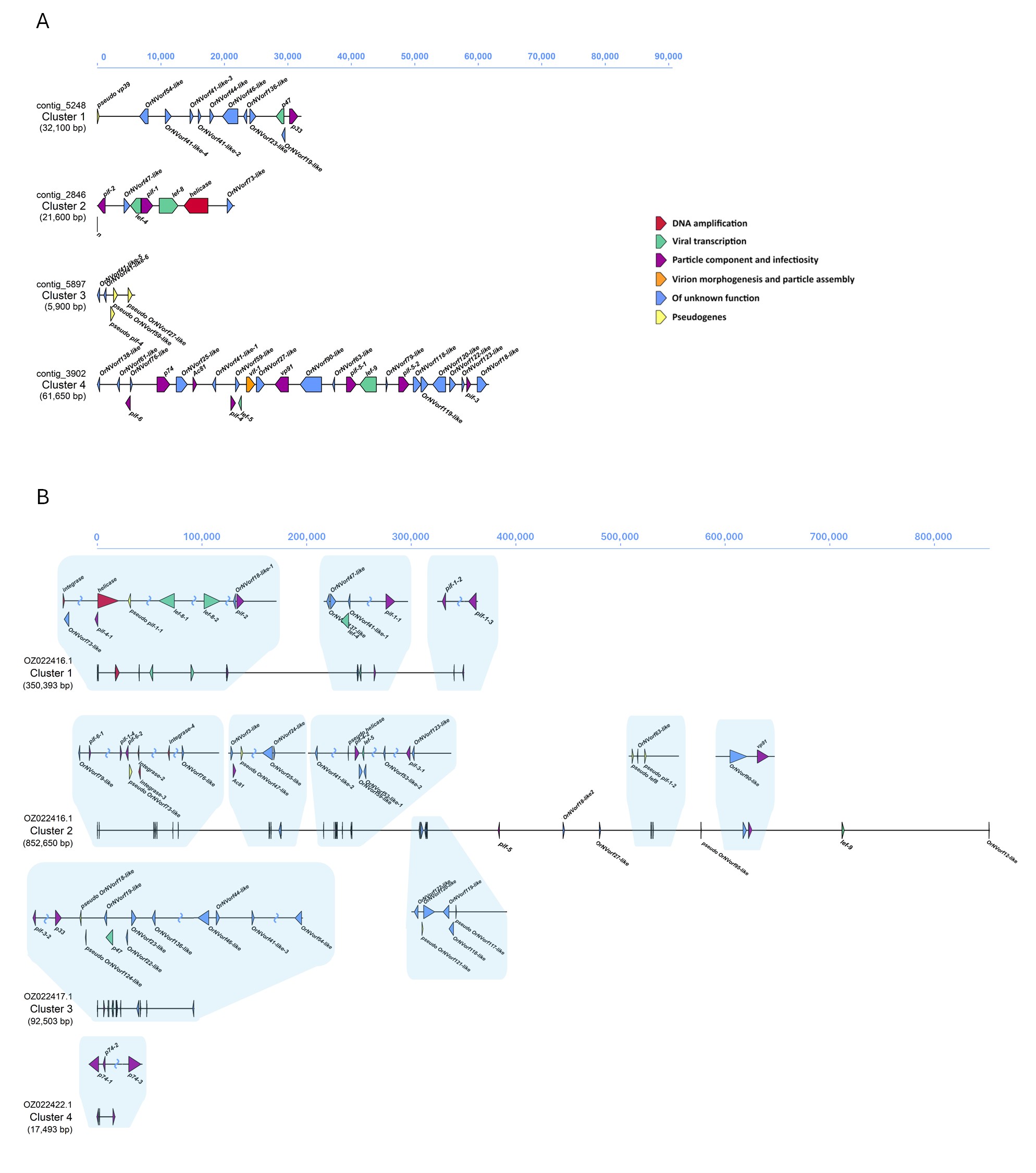

### Figure S4

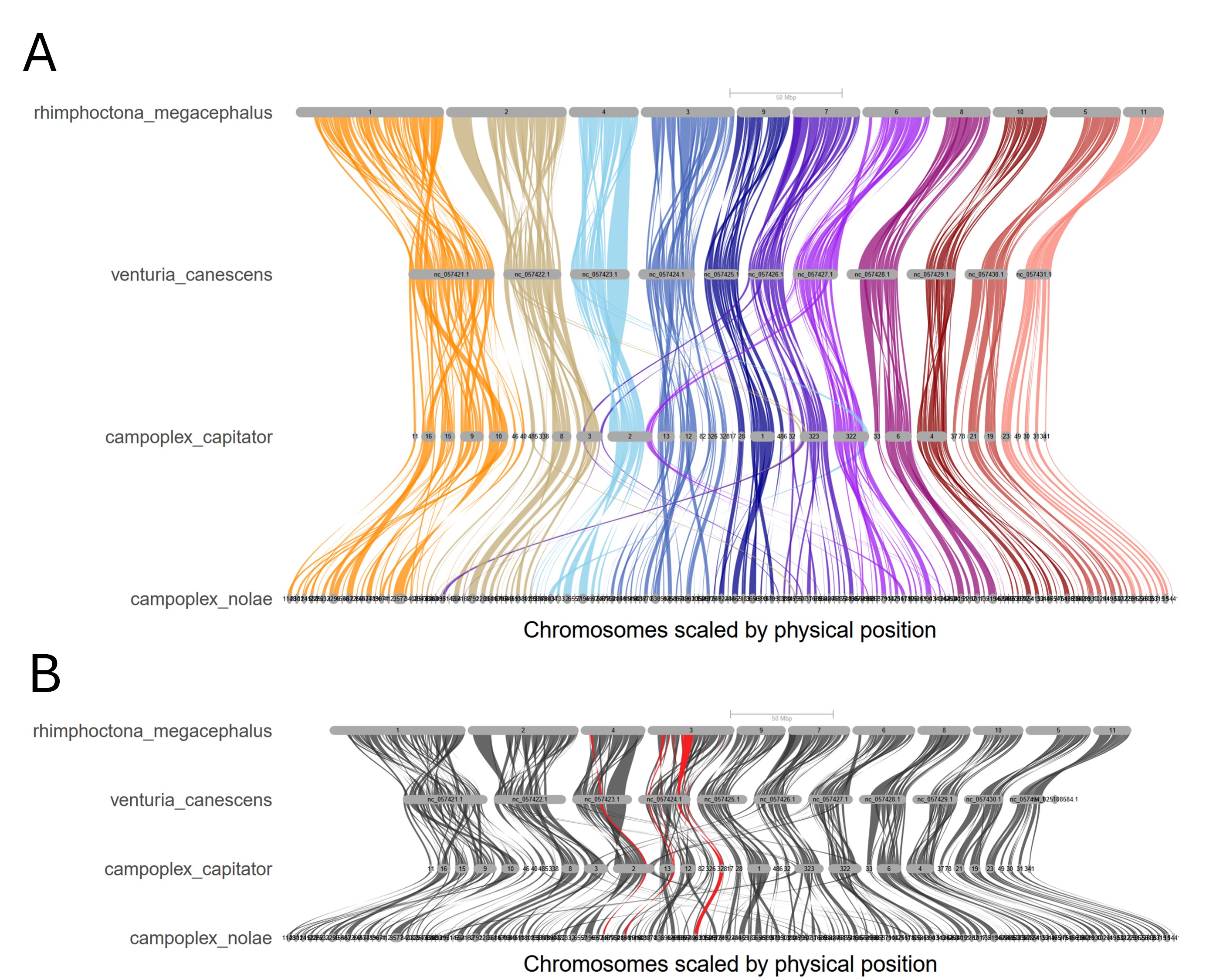

### Figure S5

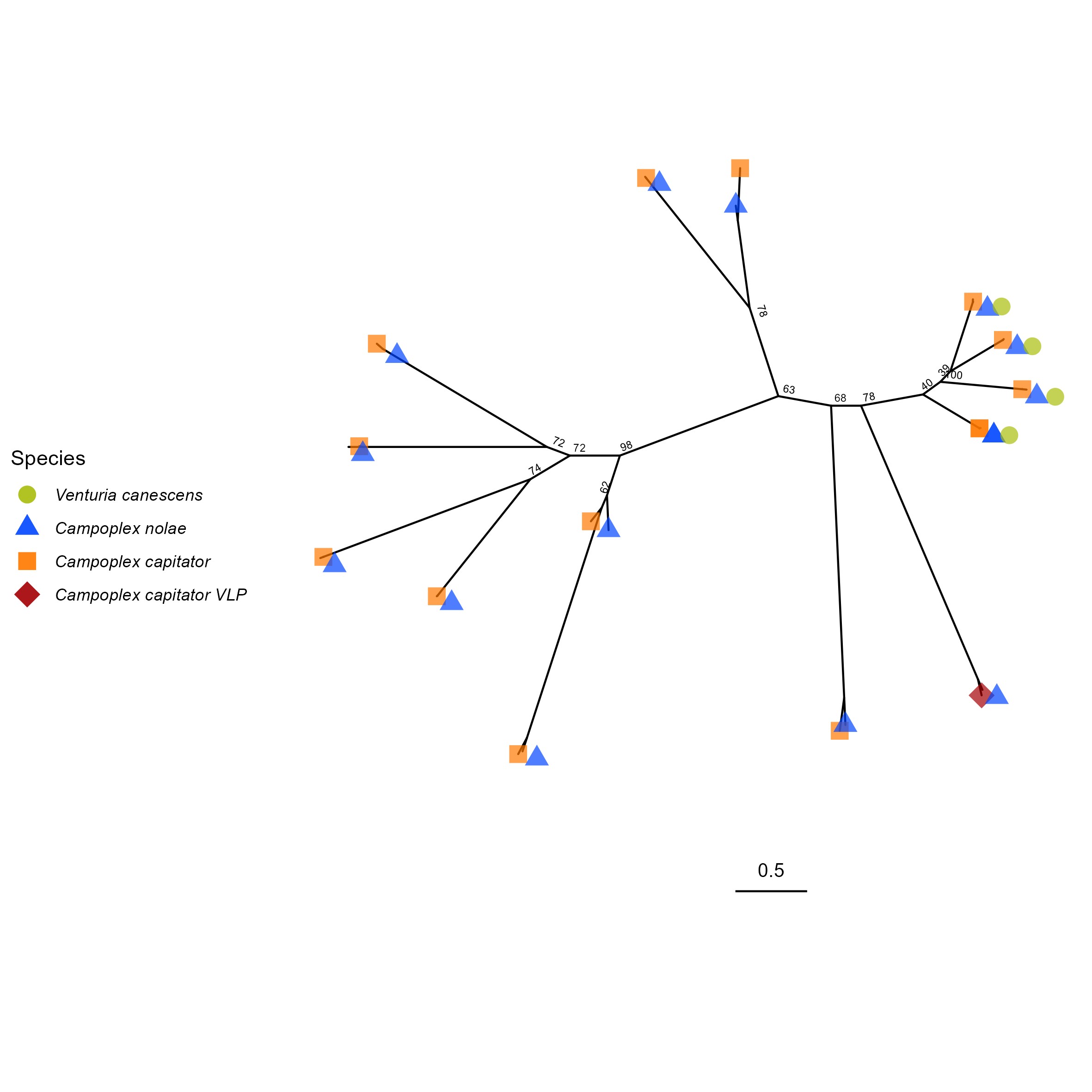
